# Susceptible host dynamics explain pathogen resilience to perturbations

**DOI:** 10.1101/2025.06.13.659551

**Authors:** Sang Woo Park, Bjarke Frost Nielsen, Emily Howerton, Bryan T. Grenfell, Sarah Cobey

## Abstract

Interventions to slow the spread of SARS-CoV-2 significantly disrupted the transmission of other pathogens. As interventions lifted, whether and when human pathogens would eventually return to their pre-pandemic dynamics remains to be answered. Here, we present a framework for estimating pathogen resilience based on how fast epidemic patterns return to their pre-pandemic dynamics. By analyzing time series data from Hong Kong, Canada, Korea, and the US, we quantify the resilience of common respiratory pathogens and further predict when each pathogen will eventually return to its pre-pandemic dynamics. Our predictions are able to distinguish which pathogens should have returned already, and deviations from these predictions reveal long-term impacts of pandemic perturbations. We find a faster rate of susceptible replenishment underlies pathogen resilience and sensitivity to both large and small perturbations. Overall, our analysis highlights the persistent nature of common respiratory pathogens compared to vaccine-preventable infections, such as measles.

**Significance Statement:** COVID-19 interventions slowed the transmission of many respiratory pathogens in different ways, raising questions about the mechanisms driving the variation in responses to interventions. To address this gap, we characterized the sensitivity of pathogen transmission to perturbations by quantifying how fast each pathogen returned to its pre-pandemic circulation patterns. We analyzed data from Hong Kong, Canada, Korea, and the US, and showed that common respiratory pathogens are far less sensitive to perturbations than measles, a vaccine-preventable infection. Finally, we showed that the speed of replenishment of the susceptible population—for example, through waning immunity—largely determines this sensitivity, suggesting that the persistence of common respiratory pathogens is likely driven by rapid susceptible replenishment.

## Introduction

Non-pharmaceutical interventions to slow the spread of SARS-CoV-2 disrupted the transmission of other human respiratory pathogens, adding uncertainties to their future epidemic dynamics and their public health burden [1]. As interventions lifted, large differences in outbreak dynamics were observed across different pathogens in different countries, with some pathogens exhibiting earlier and faster resurgences than others [2, 3, 4]. These differences likely reflect variation in interventions, pathogen characteristics, immigration/importation from other countries, and pre-pandemic pathogen dynamics [5]. Therefore, comparing the differential impacts of pandemic perturbations can provide unique opportunities to learn about underlying pathogen characteristics in different populations, such as pathogens’ transmissibility and duration of immunity [6].

Even though more than five years have passed since the emergence of SARS-CoV-2, we still observe persistent changes in pathogen dynamics following the pandemic perturbations. For example, compared to pre-pandemic, seasonal patterns, human metapneumovirus in Korea seems to circulate at lower levels, whereas RSV in Korea seems to exhibit different seasonality (Figure 1). These observations suggest the possibility of a long-term change in pathogen dynamics following the pandemic perturbations, which might be driven by a long-term shift in human behavior or population-level immunity [7, 8]. For example, the emergence of SARS-CoV-2 could have caused a long-term shift in population-level immunity through interactions with other pathogens [9], especially seasonal coronaviruses [7, 10, 11]. The possibility of a long-lasting impact of the pandemic perturbations poses an important question about future outbreaks: can we predict whether and when other pathogens will eventually return to their pre-pandemic circulation patterns?

**Figure 1:**
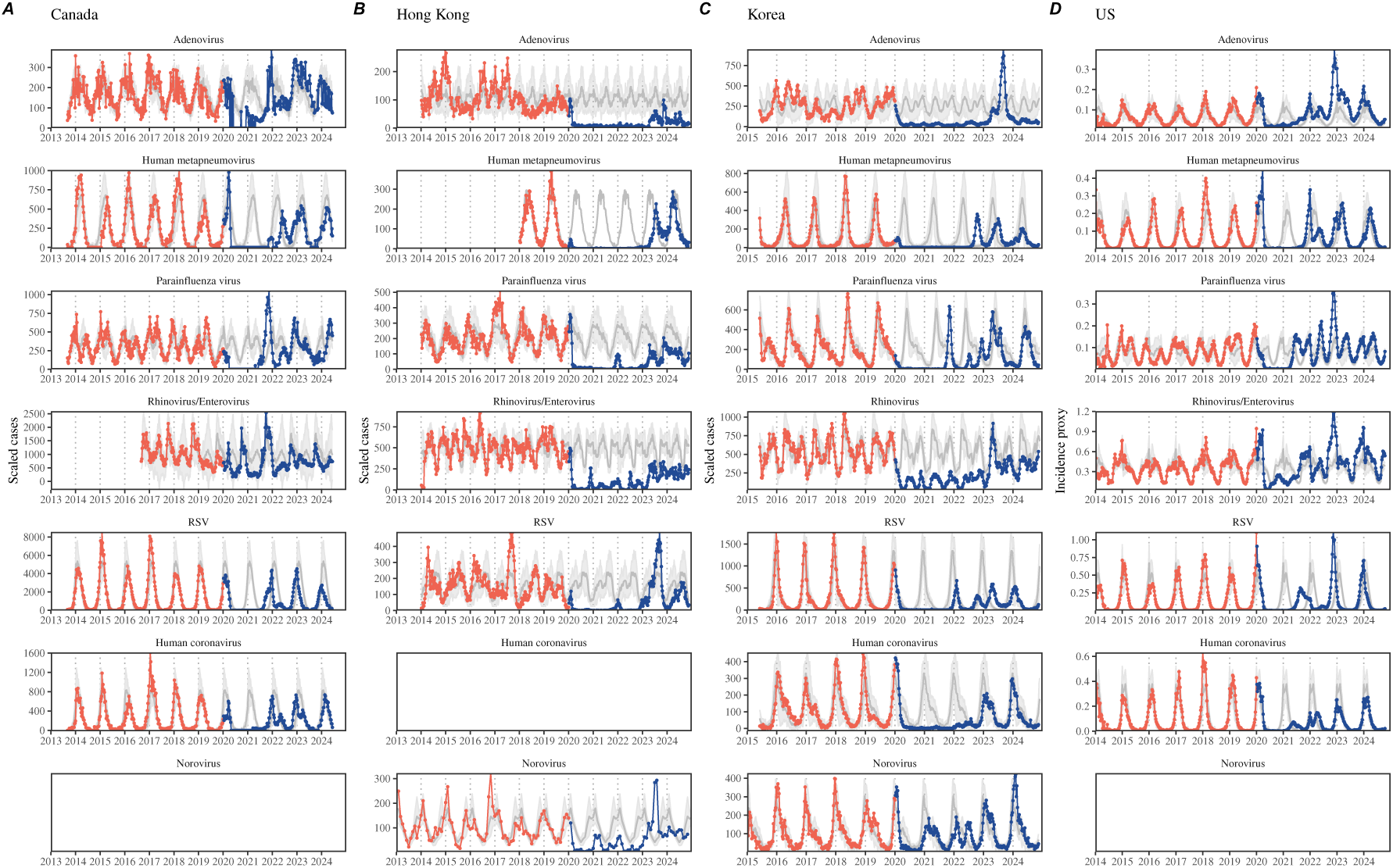
Observed heterogeneity in responses to pandemic perturbations across respiratory pathogens and norovirus in (A) Canada, (B) Hong Kong, (C) Korea, and (D) US. Red points and lines represent data before 2020. Blue points and lines represent data since 2020. Gray lines and shaded regions represent the mean seasonal patterns and corresponding 95% confidence intervals around the mean. Mean seasonal patterns were calculated by aggregating cases before 2020 into 52 weekly bins and taking the average in each week. Cases were scaled to account for changes in testing patterns (Materials and Methods).

So far, most analyses of respiratory pathogens after pandemic perturbations have focused on characterizing the timing of rebound (i.e., when the incidence first returns above historical thresholds) and linking this timing to the relaxation of intervention measures or changes in population susceptibility [1, 12, 5]. While quantifying this timing is useful for public health agencies, epidemic rebound differs from the return to historical dynamics—the rebound marks the initial return of pathogen activity without any information about the subsequent dynamics. Instead, we seek to characterize how fast (and whether) a pathogen returns to its pre-pandemic dynamics by utilizing dynamical information after the perturbation. Measuring this rate of return is useful because it allows us to quantify the ecological resilience of a host-pathogen system, which can inform responses to future interventions [13, 14, 15, 16].

In this study, we explore theoretical and statistical approaches to characterizing the resilience of a host-pathogen system based on how fast the system recovers from perturbation. We begin by laying out representative scenarios that capture the potential impacts of pandemic perturbations on endemic pathogen dynamics and illustrate how resilience can be measured by comparing the pre- and post-pandemic dynamics of susceptible and infected hosts. In practice, information on susceptible hosts is often unavailable, making this theoretical approach infeasible. Instead, we utilize a mathematical technique to reconstruct attractors from the data [17], which allows us to measure the rate at which the host-pathogen system approaches this empirical attractor after a perturbation; we define this rate to be the empirical resilience of the host-pathogen system. We use this method to analyze pathogen surveillance data for respiratory and non-respiratory pathogens from Canada, Hong Kong, Korea, and the US. Finally, we show that susceptible host dynamics explain variation in pathogen resilience and demonstrate that more resilient pathogens will be less sensitive to perturbations caused by demographic stochasticity, providing a direct link between pathogen resilience and persistence.

### Conceptual introduction to pathogen resilience

In the classical ecological literature, the resilience of an ecological system is measured by the rate at which the system is expected to return to its reference state following a perturbation [13, 14, 15, 16]. This rate corresponds to the largest real part of the eigenvalues of the linearized system near equilibrium—here, we refer to this value as the *intrinsic* resilience of the system. In practice, we rarely know the true, underlying data-generating process that describes the dynamics of common respiratory pathogens, which can depend on hidden variables, such as changes in population-level susceptibility. Instead, we have to rely on simplifying model assumptions, raising questions about whether we can infer the intrinsic resilience of a system accurately. Nonetheless, we can measure the *empirical* resilience of a host-pathogen system by looking at how fast the system returns to the pre-perturbation endemic dynamics after the perturbation has ended. The COVID-19 pandemic provides a convenient example of a major perturbation.

### Resilience of a single-strain system under a short-term perturbation

As an example, we begin with a simple Susceptible-Infected-Recovered-Susceptible (SIRS) model with seasonally forced transmission and demography (i.e., birth and death) [18]. Here, a pandemic perturbation that reduces transmission by 50% for 6 months starting in 2020 causes the epidemic to deviate from its original stable annual cycle and eventually return (Figure 2A). To quantify resilience, we measure the distance from the pre-pandemic attractor—a set of points in state space or phase plane that the system is pulled towards [19]. Specifically, we measure the Euclidean distance on the SI phase plane relative to the counterfactual unperturbed trajectory (Figure 2B; Materials and Methods). This distance decreases exponentially with time on average, as demonstrated by the locally estimated scatterplot smoothing (LOESS) fit, and the overall rate of return matches the analytically derived intrinsic resilience of the seasonally unforced system (Figure 2C).

**Figure 2:**
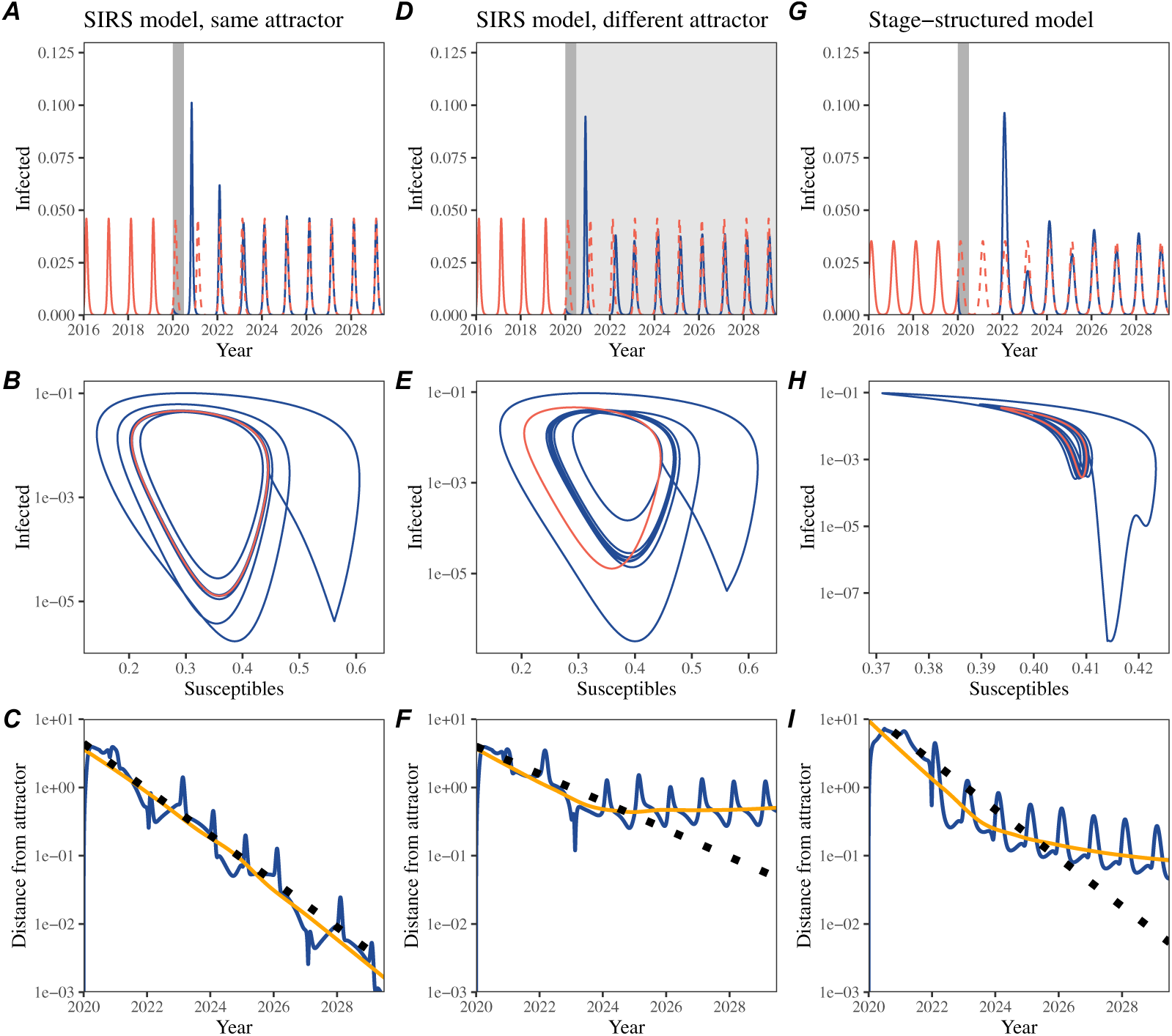
A simple method to measure pathogen resilience following pandemic perturbations across different scenarios. (A, D, G) Simulated epidemic trajectories across various models. Red and blue solid lines represent epidemic dynamics before and after pandemic perturbations are introduced, respectively. Red dashed lines represent counterfactual epidemic dynamics in the absence of perturbations. Gray regions indicate the duration of perturbations with the dark gray region representing a 50% transmission reduction and the light gray region representing a 10% transmission reduction. (B, E, H) Phase plane representation of the time series in panels A, D, and G alongside the corresponding susceptible host dynamics. Red and blue solid lines represent epidemic trajectories on an SI phase plane before and after perturbations are introduced, respectively. (C, F, I) Changes in distance from the attractor over time on a log scale. Blue lines represent the distance from the attractor. Orange lines represent the locally estimated scatterplot smoothing (LOESS) fits to the logged distance from the attractor. Dotted lines show the intrinsic resilience of the seasonally unforced system.

### Resilience of a single-strain system under a long-term perturbation

Alternatively, pandemic perturbations can have a lasting impact on pathogen dynamics through a reduction in transmission or change in immunity. For example, if a 10% reduction in transmission persists even after the major pandemic perturbations are lifted, the system will converge to a new attractor (Figure 2D–F), but we may not be able to confirm the convergence to a new attractor until many years have passed. Nonetheless, we can still measure the distance from the pre-pandemic attractor (Figure 2E). The distance initially decreases exponentially on average (equivalently, linearly on a log scale) and eventually plateaus (Figure 2F). Here, a permanent 10% reduction in transmission rate slows the system, causing a slower decrease in the distance from the attractor (Figure 2F) than it would have otherwise (Figure 2C). This example shows that resilience is not necessarily an intrinsic property of a specific pathogen. Instead, pathogen resilience is a property of a specific attractor that a host-pathogen system approaches, which depends on both pathogen and host characteristics.

### Resilience of a single-strain system with long-term transients

Finally, transient phenomena can further complicate the picture (Figure 2G–I). For example, a stage-structured model that includes a reduction in secondary susceptibility initially exhibits a stable annual cycle, but perturbations from a 50% reduction in transmission for 6 months cause the epidemic to shift to a transient biennial cycle before it eventually approaches the original pre-pandemic attractor (Figure 2G,H). As indicated by the LOESS fit, the distance from the attractor initially decreases exponentially at a rate that is consistent with the intrinsic resilience of the seasonally unforced stage-structured system, but the rate of return slows with the damped oscillations (Figure 2I). This behavior is also referred to as a ghost attractor, which causes long transient dynamics and slow transitions [20]. Strong seasonal forcing in transmission can also lead to transient phenomena for a simple SIRS model, causing a slow return to pre-perturbation dynamics (Supplementary Figure S1). However, adding a small rate of imported infections to this strongly seasonal model removes these transient dynamics and increases the system’s resilience (Supplementary Figure S2). Such a change to a non-seasonal model has considerably reduced impact on resilience (Supplementary Figure S3).

### Resilience of a two-strain system

This empirical approach allows us to measure the resilience of a two-strain host-pathogen system even when we imperfectly observe infection dynamics. Simulations of two competing strains illustrate that separate analyses of individual strain dynamics (e.g., RSV subtype A vs B) and a joint analysis of total infections (e.g., total RSV infections) yield identical resilience estimates (Supplementary Figure S4, 5). This is expected because eigenvalues determine the dynamics of the entire system around the equilibrium, meaning that both strains should exhibit identical rates of return following a perturbation. Analogous to a single-strain system, strong seasonal forcing in transmission can cause the two-strain system to slow down through transient phenomena (Supplementary Figure S6).

These observations yield three insights. First, we can directly estimate the empirical resilience of a host-pathogen system by measuring the rate at which the system approaches an attractor. The empirical approach to estimating pathogen resilience is particularly convenient because it does not require us to know the true, underlying data-generating process; estimating the intrinsic resilience from fitting misspecified models can lead to biased estimates (Supplementary Figure S7). Second, resilience estimates allow us to make phenomenological predictions about the dynamics of a host-pathogen system following a perturbation. Assuming that an attractor has not changed and the distance from the attractor will decrease exponentially over time, we can estimate when the system should reach an attractor. Finally, a change in the (exponential) rate of approach can provide information about whether the system has reached an alternative attractor, or a ghost attractor, that is different from the original, pre-pandemic attractor. These alternative attractors may reflect permanent changes in transmission patterns as well as changes in immune landscapes. Now that several years have passed since major interventions have been lifted, many respiratory pathogens may have had sufficient time to return to their post-intervention attractors—empirical comparisons between current cases and pre-pandemic cases are consistent with this hypothesis (Figure 1). With recent data, we can start to evaluate whether we see early signs of convergence to the former attractor or a new one.

### Inferring pathogen resilience from real data

Based on these patterns, we now present our approach to estimating pathogen resilience from real data (Figure 3). We first tested this approach against simulations and then analyzed case time series of respiratory pathogens from Canada, Hong Kong, Korea, and the US.

**Figure 3:**
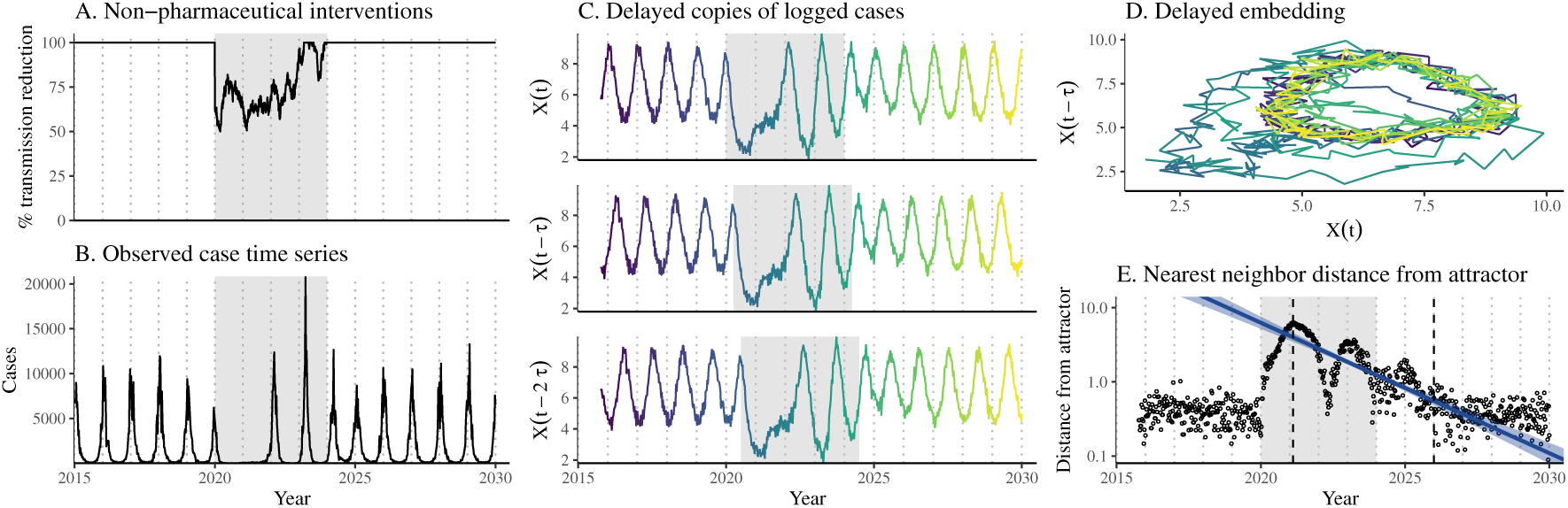
A schematic diagram explaining the inference of pathogen resilience from synthetic data. (A) A realistic example of simulated pandemic perturbations, represented by a relative reduction in transmission. (B) The impact of the synthetic pandemic perturbation on epidemic dynamics simulated using a SIRS model with demographic stochasticity. (C) Generating delayed copies of the logged time series allows us to obtain an embedding. (D) Two dimensional representation of an embedding. (E) Delayed embedding allows us to calculate the nearest neighbor distance from the empirical attractor, which is determined based on the pre-pandemic time series. Blue lines and dashed regions represent the linear regression fit and the associated 95% confidence interval. This distance time series can be used to infer pathogen resilience after choosing an appropriate window for linear regression. For illustration purposes, an arbitrary fitting window was chosen.

So far, we have focused on simple examples that assume a constant transmission reduction during the pandemic. However, in practice, the impact of pandemic perturbations on pathogen transmission is likely more complex (Figure 3A), reflecting the introduction and relaxation of various intervention strategies. For example, continued transmission reduction from interventions can limit the size of the first outbreak following the rebound, allowing for a larger outbreak when interventions are further relaxed (Figure 3B).

Previously, we relied on the dynamics of susceptible and infected hosts to compute the distance from the attractor (Figure 2), but information on susceptible hosts is rarely available in practice. In addition, uncertainties in case counts due to observation error, strain evolution, and multiannual cycles in the observed epidemic dynamics (e.g., adenovirus in Hong Kong and Korea) complicate defining pre-pandemic attractors. To address these challenges, we can reconstruct an empirical attractor by utilizing Takens’ theorem [17], which states that an attractor of a nonlinear multidimensional system can be mapped onto a delayed embedding (Materials and Methods). Specifically, by taking the logged values of pre-pandemic cases log(*C*(*t*) + 1) and creating delayed copies, Takens’ theorem allows us to define the pre-pandemic attractor as a set of points in an *M* -dimensional space:

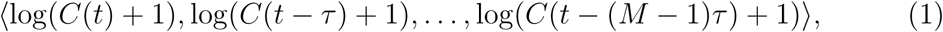

where the delay *τ* and embedding dimension *M* are determined based on autocorrelations and false nearest neighbors, respectively [21, 22]. Intuitively, Takens’ theorem tells us that, delayed copies of the observed time series preserve the dynamical properties of the underlying multi-dimensional attractor. We can then apply the same delay and embedding dimensions to the entire time series to determine its position in multi-dimensional state space (Figure 3D) and measure the nearest neighbor distance between the current state of the system and the empirical pre-pandemic attractor (Figure 3E).

In theory, the rate at which the distance from this attractor decreases, which can be estimated by fitting a linear regression on a log scale, provides an empirical measure of pathogen resilience (Figure 2C,F,I; Figure 3E). However, resulting estimates of pathogen resilience can be sensitive to choices about embedding delays and dimensions. Longer delays and higher dimensions tend to smooth out temporal variations in the distance from the attractor (Supplementary Figure S8).

Estimating pathogen resilience from linear regression is also sensitive to the choice of window used to fit the regression (Figure 3E). Therefore, we validated our criteria for window selection and embedding dimensions using simulations (Materials and Methods; Supplementary Results; Supplementary Figure S9). We then applied these criteria to analyze pathogen surveillance data presented in Figure 1 (Materials and Methods; Supplementary Results; Supplementary Figure S10-S11).

For all pathogens we considered, resilience estimates fall between 0.4/year and 1.8/year (Figure 4A). We estimated the mean resilience of common respiratory pathogens to be 0.99/year (95% CI: 0.81/year–1.18/year). For reference, this is ≈ 7.5 times higher than the intrinsic resilience of pre-vaccination measles in England and Wales (≈ 0.13/year). Finally, resilience estimates for norovirus, a gastrointestinal pathogen, were comparable to those of common respiratory pathogens: 1.44/year (95% CI: 1.01/year–1.87/year) for Hong Kong and 1.07/year (95% CI: 0.86/year– 1.29/year) for Korea. Based on a simple ANOVA test, we did not find significant differences in resilience estimates across countries (*p* = 0.25) or pathogens (*p* = 0.67). Using resilience estimates, we predicted when each pathogen would hypothetically return to its pre-pandemic dynamics, assuming no long-term change in the attractor.

**Figure 4:**
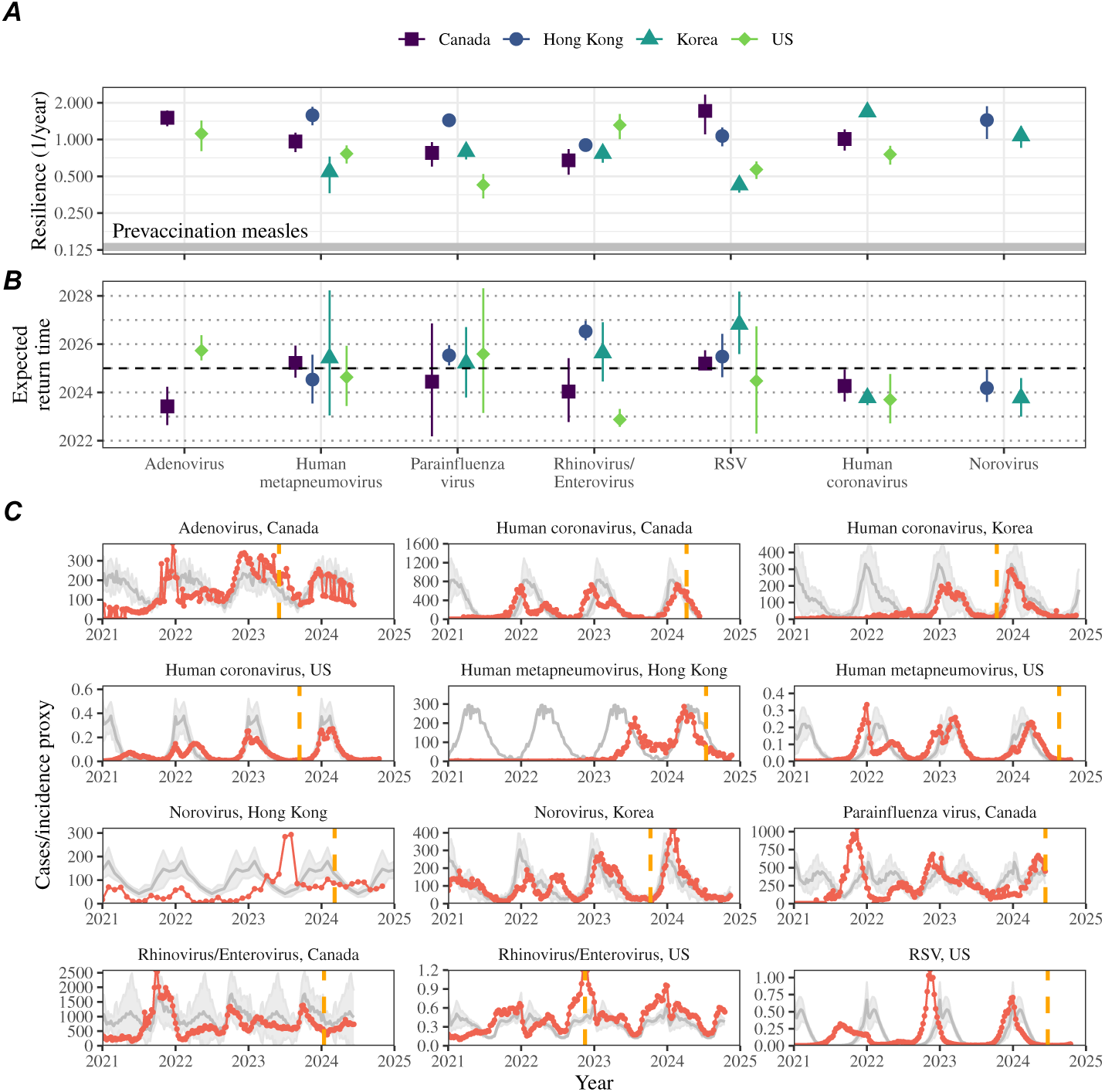
Summary of resilience estimates and predictions for return time. (A) Estimated pathogen resilience. The gray horizontal line represents the intrinsic resilience of pre-vaccination measles dynamics. (B) Predicted timing of when each pathogen will return to their pre-pandemic cycles. The dashed line in panel B indicates the end of 2024. Error bars represent 95% confidence intervals. (C) Observed dynamics for pathogen that are predicted to have returned before the end of 2024. Red points and lines represent data since 2020. Gray lines and shaded regions represent the mean seasonal patterns and corresponding 95% confidence intervals around the mean, previously shown in Figure 1. Orange vertical lines represent the predicted timing of return.

Specifically, we extended our linear regression fits to distance-from-attractor time series and ask when the predicted regression line will cross a threshold value; since we relied on nearest neighbor distances, pre-pandemic distances are always greater than zero (Figure 3E), meaning that we can use the mean of pre-pandemic distances as our threshold.

We predicted that a return to pre-pandemic cycles has occurred or would be imminent for most pathogens (Figure 4B). In particular, we predicted that 12 out of 23 pathogen-country pairs should have already returned before the end of 2024. For almost all pathogens that were predicted to have returned already, the observed epidemic dynamics showed clear convergence towards their pre-pandemic seasonal averages, confirming our predictions (Figure 4C). However, there were a few exceptions, including norovirus in Hong Kong and rhinovirus/enterovirus in the US, where the observed epidemic dynamics in 2024 exhibit clear deviation from their prepandemic seasonal averages (Figure 4C; Figure S11). These observations suggest a possibility that some common respiratory pathogens may have converged to different attractors or are still exhibiting non-equilibrium dynamics. In contrast, pathogens that were predicted to have not returned also showed clear differences from their pre-pandemic seasonal averages; as many of these pathogens are predicted to return in 2025–2026, we may be able to test these predictions in near future (Supplementary Figure S12). Our reconstructions of distance time series and estimates of pathogen resilience and expected return time were generally robust to choices of embedding dimensions (Supplementary Figure S13–14). We found that making out-of-sample predictions were challenging, but still supported consistency in our predictions (Supplementary Results; Supplementary Figure S15–16).

### Susceptible host dynamics explain variation in pathogen resilience

So far, we have focused on quantifying pathogen resilience from the observed patterns of pathogen re-emergence following pandemic perturbations. But what factors determine the resilience of a host-pathogen system? Here, we illustrate how susceptible host dynamics affect pathogen resilience by varying the basic reproduction number R_0_ and the duration of immunity of the SIRS model.

An increase in R_0_ and a decrease in the duration of immunity increase resilience (Figure 5A) and the per-capita rate of susceptible replenishment, the ratio between the total rate at which new susceptibles enter the population and the equilibrium number of susceptibles, *S*^∗^ (Figure 5B). Specifically, a higher R_0_ drives a faster percapita susceptible replenishment rate by decreasing *S*^∗^. For a simple SIR model that assumes life-long immunity, pathogen resilience is proportional to the per-capita rate of susceptible replenishment (Materials and Methods). Overall, these observations suggest that a faster per-capita susceptible replenishment rate causes the system to be more resilient.

**Figure 5:**
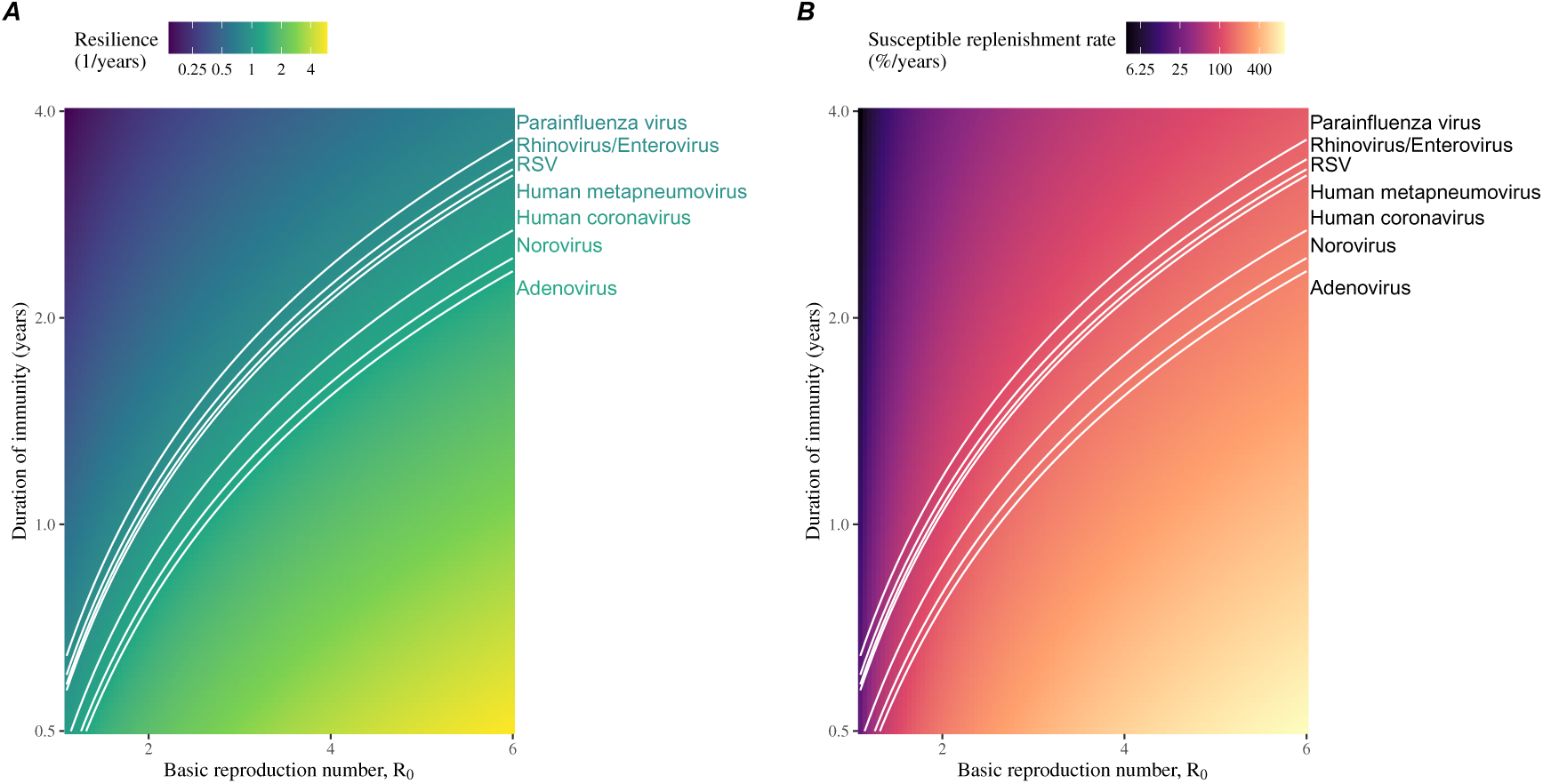
Linking pathogen resilience to epidemiological parameters and susceptible host dynamics. (A) Intrinsic resilience as a function of the basic reproduction number R_0_ and the duration of immunity. (B) Per-capita susceptible replenishment rate as a function of the basic reproduction number R_0_ and the duration of immunity. The standard SIRS model without seasonal forcing is used to compute intrinsic resilience and the per-capita susceptible replenishment rate. Lines correspond to a set of parameters that are consistent with mean resilience estimates for each pathogen, averaged across different countries.

By taking the average values of empirical resilience for each pathogen, we were able to map each pathogen onto a set of parameters of the SIRS model that are consistent with corresponding resilience estimates (Figure 5A). Across all pathogens we considered, we estimated that the average duration of immunity is likely to be short (*<* 4 years) across a plausible range of R_0_ (*<* 6). In contrast, there was large uncertainty in the estimates for susceptible replenishment rates due to a lack of one-to-one correspondence between susceptible replenishment rates and pathogen resilience (Figure 5B).

### Pathogen resilience determines sensitivity to stochastic perturbations

Even in the absence of major pandemics, host-pathogen systems are expected to experience continued perturbations of varying degrees from changes in epidemiological conditions, such as human behavior, climate, and viral evolution. These perturbations can also arise from demographic stochasticity, which is inherent to any ecological system. Here, a seasonally unforced SIRS model with constant birth and death rates illustrates that resilience determines the sensitivity to perturbations caused by demographic stochasticity (Figure 6, Materials and Methods). Less resilient systems exhibit greater deviations from the deterministic trajectory and slower epidemic cycles (Figure 6A–C). The periodicity of this epidemic cycle matches those predicted by the intrinsic periodicity of the system (Supplementary Figure S17), which is inversely proportional to the intrinsic resilience (Supplementary Figure S18).

**Figure 6:**
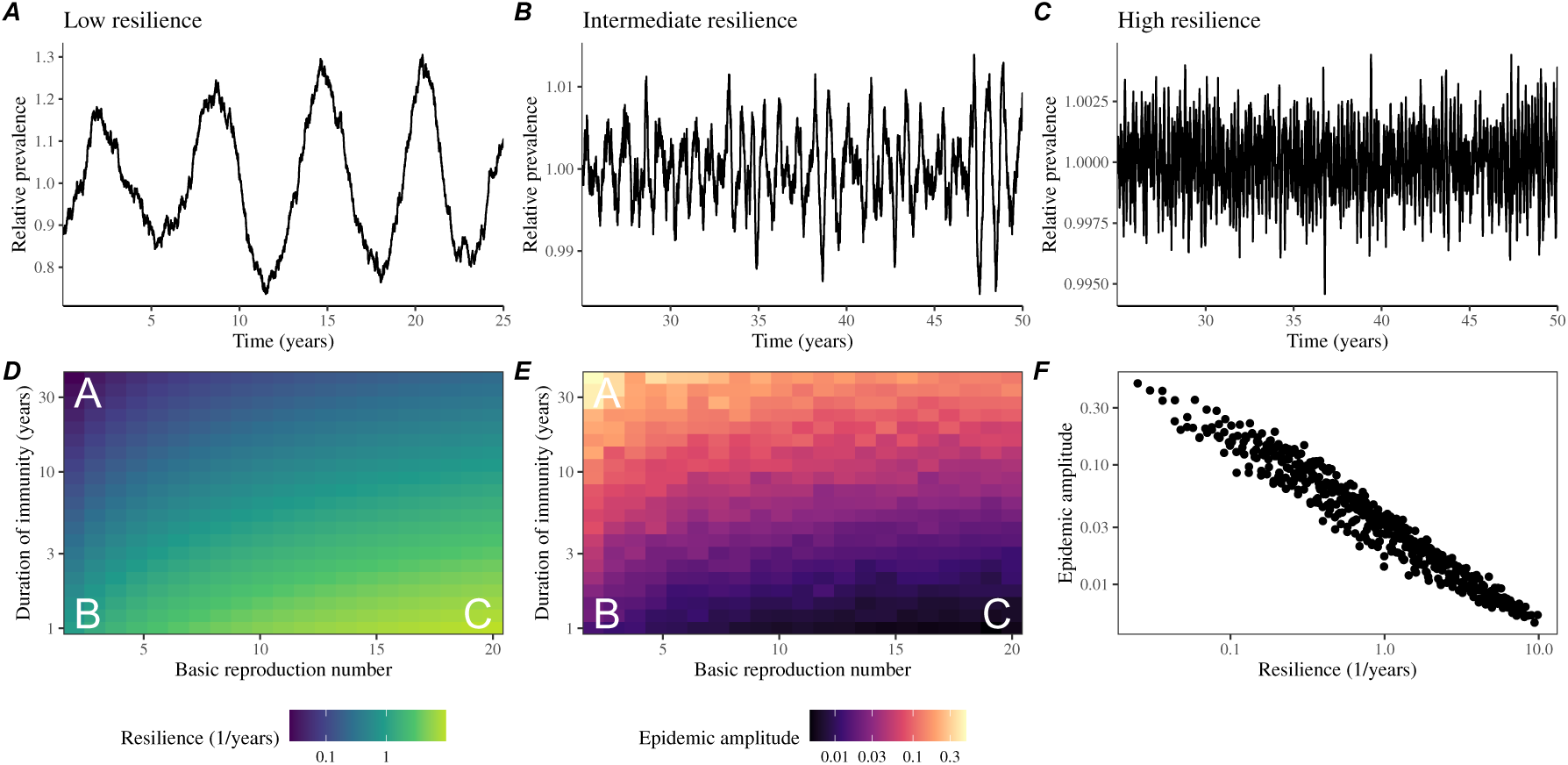
Linking resilience of a host-pathogen system to its sensitivity to stochastic perturbations. (A–C) Epidemic trajectories of a stochastic SIRS model across three different resilience values: low, intermediate, and high. The relative prevalence was calculated by dividing infection prevalence by its mean value. (D) Intrinsic resilience of a system as a function of the basic reproduction number R_0_ and the duration of immunity. (E) Epidemic amplitude as a function of the basic reproduction number R_0_ and the duration of immunity. The epidemic amplitude corresponds to (max *I* −min *I*)/(2*Ī*), where *Ī* represents the mean prevalence. Labels A–C in panels D and E correspond to scenarios shown in panels A–C. (F) The relationship between pathogen resilience and epidemic amplitude.

The intrinsic resilience is not the sole determinant of a system’s sensitivity to stochastic perturbations. For example, the population size and average duration of infection also affect the deviation from the deterministic trajectory caused by demographic stochasticity, even though these quantities have little to no impact on the intrinsic resilience (Supplementary Figure S17). These conclusions were robust for the seasonally forced SIRS model, where the amount of deviation from the deterministic trajectory depends on the corresponding resilience of the seasonally unforced system (Supplementary Figure S19).

## Discussion

COVID-19 pandemic interventions caused major disruptions to circulation patterns of respiratory and non-respiratory pathogens, adding challenges to predicting their future dynamics [1, 2, 3, 4]. These perturbations offer large-scale natural experiments for understanding how different pathogens respond to perturbations. In this study, we showed that pathogen re-emergence patterns following pandemic perturbations can be characterized through the lens of ecological resilience, and we presented a new method for estimating pathogen resilience from time series data. We showed that variation in pathogen resilience can be explained by the differences in susceptible host dynamics, where faster replenishment of the susceptible pool corresponds to a more resilient host-pathogen system. Finally, we showed that pathogen resilience also determines the sensitivity to stochastic perturbations. The link between responses to large pandemic perturbations and small stochastic perturbations via pathogen resilience echoes fluctuation–response principles observed in other biological systems [23].

We analyzed case time series of common respiratory infections and norovirus infections from Canada, Hong Kong, Korea, and the US to estimate their resilience. Overall, we estimated the resilience of these pathogens to range from 0.4/year to 1.8/year, which is 7.5 times more resilient, on average, than prevaccination measles. Consistent with other epidemiological evidence [24, 25, 26, 27], these resilience estimates indicate that common respiratory pathogens and norovirus likely exhibit faster susceptible replenishment and are therefore more persistent, indicating potential challenges in controlling these pathogens.

Based on our resilience estimates, we made phenomenological predictions about when each pathogen will return to its endemic cycle. Our main regression analysis could accurately distinguish which pathogens should have already returned before the end of 2024. However, there were two main exceptions (i.e., norovirus in Hong Kong and rhinovirus/enterovirus in the US), hinting at long-lasting impact of pandemic perturbations. These changes may reflect changes in surveillance or the dynamics. While it may seem unlikely that behavioral changes would only affect a few pathogens and not others, we cannot rule out this possibility given differences in the observed mean age of infections and therefore the differences in age groups that primarily drive transmission [28, 29]. Differences in the mode of transmission between respiratory and gastrointestinal pathogens may also contribute to divergent responses to pandemic perturbations.

For almost half of the pathogens we considered, we predicted that their return to original epidemic patterns is imminent. However, our evaluation of out-of-sample predictions suggested major challenges in predicting the exact timing of pathogen return and estimating pathogen resilience from limited data. Therefore, our estimates of prediction timing should be taken as a ballpark estimate. We will need a few more years of data to test whether these pathogens will eventually return to their original dynamics or eventually converge to a different attractor. We also cannot rule out the possibility that some pathogens may exhibit long-term transient behaviors following pandemic perturbations. Overall, these observations echo earlier studies that highlighted the long-lasting impact of pandemic perturbations [8, 30, 31, 4, 32]. We showed that susceptible host dynamics shape pathogen resilience, where faster replenishment of the susceptible population causes the pathogen to be more resilient. For simplicity, we focused on waning immunity and birth as the main drivers of the susceptible host dynamics but other mechanisms can contribute to the replenishment of the susceptible population. Pathogen evolution, especially the emergence of antigenically novel strains, can cause effective waning of immunity in the population; therefore, we hypothesize that the rate of antigenic evolution is likely a key feature of pathogen resilience. Future studies should explore the relationship between the rate of evolution and resilience for antigenically evolving pathogens. This result also highlights the importance of detailed measurements of changes in the susceptible population through immune assays for understanding pathogen dynamics [33].

Many other factors may affect the resilience of a host-pathogen system beyond the susceptible replenishment rate. For example, we showed that importation can also increase the resilience of the system (Supplementary Figures S2, 3). Interestingly, this effect is minimal without seasonal forcing but amplified under strong seasonal forcing, suggesting that analyzing the eigenvalues of the seasonally unforced system may be insufficient to capture key aspects of the system’s resilience. Future studies should consider both analytical and numerical analyses of more detailed, mechanistic models to understand how other factors—such as the generation-interval distribution [34], immune boosting [35], strain competition [36], and post-infection mortality [37]—may alter resilience. Quantifying relative changes in pathogen resilience with respect to underlying parameters of a model—analogous to sensitivity and elasticity analyses in ecology [38, 39]—may allow one to infer relative importance of different factors that affect pathogen resilience.

There are several limitations to our work. First, we did not extensively explore other approaches to reconstructing the attractor. Recent studies showed that more sophisticated approaches, such as using non-uniform embedding, can provide more robust reconstruction for noisy data [22]. In the context of causal inference, choices about embedding can have big impacts on the resulting inference [40]. Our resilience estimates are likely overly confident given a lack of uncertainties in attractor reconstruction as well as the simplicity of our statistical framework. Nonetheless, as illustrated in our sensitivity analyses, inferences about pathogen resilience in our SIRS model appear to be robust to decisions about embedding lags and dimensions. Short pre-pandemic time series also limit our ability to accurately reconstruct the attractor and contribute to the crudeness of our resilience estimates. Our framework also does not allow us to distinguish whether a system has settled to a new attractor or is experiencing long-term transient behavior.

Uncertainties in pathogen dynamics due to changes in testing patterns further contribute to the crudeness of our resilience estimates. We tried to account for changes in testing patterns by either scaling the case counts (for Canada, Hong Kong, and Korea) or deriving an incidence proxy (for the US). We were unable to derive an incidence proxy for Canada and Hong Kong due to instability of the test positivity ratio. Therefore, we cannot rule out the possibility that the apparent long-term impact of pandemic perturbations may simply reflect changes in reporting, rather than actual changes in the dynamics. More stable proxies of incidence (e.g., from wastewater) might be useful in future studies.

Finally, our simulation-based analyses primarily focused on single-strain systems, but real-world pathogens can interact with other pathogens, which can result in complex dynamics [41, 42]. To address this limitation, we considered a simple model of two competing strains (via cross immunity) and showed that the resilience of a coupled system can be measured by studying the dynamics of either strain. However, this conclusion likely depends on the strength and mechanism of strain interactions. For example, ecological interference between two unrelated pathogens [41] will likely generate weaker coupling than cross-immunity between related pathogens; in the former case, we do not necessarily expect two unrelated pathogens to have the same resilience despite their ecological interference. Some pathogen strains can also exhibit positive interactions where infection by one strain can lead to an increased transmission of another competing strain. Our study nonetheless illustrates that quantifying pathogen resilience can provide novel insights into pathogen dynamics. Furthermore, our qualitative prediction that common respiratory pathogens are more resilient than prevaccination measles is also likely to be robust, given how rapidly many respiratory pathogens returned to their original cycles following pandemic perturbations.

Predicting the impact of anthropogenic changes on infectious disease dynamics is a fundamental aim. Our study illustrates that how a host-pathogen system responds to both small and large perturbations is largely predictable through the lens of ecological resilience. In particular, quantifying the resilience of a host-pathogen system offers insight into endemic pathogens’ responses to pandemic perturbations, including whether some pathogens will experience long-lasting impacts. More broadly, a detailed understanding of the determinants of pathogen resilience might improve understanding of pathogen persistence.

## Materials and Methods

### Data

We gathered time series on respiratory infections from Canada, Hong Kong, Korea, and United States (US). As a reference, we also included norovirus infections when available. In contrast to respiratory pathogens, we hypothesized gastrointestinal viruses, such as norovirus, to be differently affected by pandemic perturbations.

Weekly time series of respiratory infection cases in Canada came from a publicly available website by the Respiratory Virus Detection Surveillance System, which collects data from select laboratories across Canada [43]. Weekly time series of respiratory infection cases in Hong Kong came from a publicly available website by the Centre for Health Protection, Department of Health [44, 45]. Weekly time series of acute respiratory infection cases in Korea came from a publicly available website by the Korea Disease Control and Prevention Agency [46]. Finally, weekly time series of respiratory infection cases in the US were obtained from the National Respiratory and Enteric Virus Surveillance System (NREVSS). Readers can request the data from NREVSS at. Time series on number of tests were also available in Canada, Hong Kong, and the US, but not in Korea.

### Data processing

For all time series, we rounded every year to 52 weeks by taking the average number of cases and tests between the 52nd and 53rd week. We also rescaled all time series to account for changes in testing patterns, which were then used for the actual analysis. For Canada, an increase in testing was observed from 2013 to 2024 (Supplementary Figure S21); this increase is associated with a decrease in test positivity during the same period (Supplementary Figure S22). To account for this increase, we calculated a 2 year moving average for the number of tests for each pathogen, which we used as a proxy for testing effort. Then, we divided the smoothed testing patterns by the smoothed value at the final week such that the testing effort has a maximum of 1. We then divided weekly cases by the testing effort to obtain a scaled case time series. A similar approach was used earlier for an analysis of RSV time series in the US to account for changes in testing patterns [26].

For Hong Kong, we applied the same scaling procedure to the time series as we did for Canada. In this case, we only adjusted for testing efforts up to the end of 2019 because there was a major reduction in testing for common respiratory pathogens between 2020 and 2023 (Supplementary Figure S23). Analogous to patterns in Canada, an increase in testing volume is associated with a decrease in test positivity (Supplementary Figure S24).

For Korea, while we did not have information on testing, the reported number of respiratory infections consistently increased from 2013 to the end of 2019, which we interpreted as changes in testing patterns (Supplementary Figure S25). Since we did not have testing numbers, we used the weekly sum of all acute respiratory viral infection cases as a proxy for testing, which were further smoothed with moving average and scaled to have a maximum of 1. For Korea, we also only adjusted for testing efforts up to the end of 2019.

In the US, there has been a large increase in testing for some respiratory pathogens, especially RSV, which could not be corrected by simple scaling (Supplementary Figure S26). Instead, we derived an incidence proxy by multiplying the test positivity with influenza-like illness positivity, which was taken from https://gis.cdc.gov/ grasp/fluview/fluportaldashboard.html. This method of estimating an incidence proxy has been recently applied in the analysis of seasonal coronaviruses [7] and *Mycoplasma pneumoniae* infections [4]. Detailed assumptions and justifications are provided in [47].

### Data summary

To make qualitative comparisons between pre- and post-perturbation dynamics of respiratory pathogen circulation patterns, we calculated the mean seasonal patterns using time series of either rescaled cases or incidence proxy estimates before 2020. We did so by taking the mean value in each week across all years before 2020. Confidence intervals around the means were calculated using a simple t test.

### Estimating pathogen resilience

In order to measure pathogen resilience from surveillance data, we first reconstructed the empirical pre-pandemic attractor of the system using Takens’ embedding theorem [17]. Specifically, for a given pathogen, we took the pre-pandemic (before 2020) case time series *C*(*t*) and reconstructed the attractor using delayed embedding with a uniform delay of *τ* and dimension *M* :

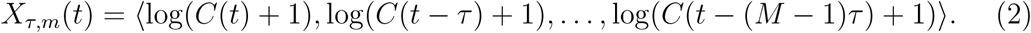

Here, the delay *τ* was determined by calculating the autocorrelation of the logged pre-pandemic time series and asking when the autocorrelation crosses 0 for the first time [22]; a typical delay for an annual outbreak is around 13 weeks.

Then, for a given delay *τ*, we determined the embedding dimension *M* using the false nearest neighbors approach [21, 22]. To do so, we started with an embedding dimension *e* and constructed a set of points *A_τ,e_* = {*X_τ,e_*(*t*)|*t <* 2020}. Then, for each point *X_τ,e_*(*t*), we determined the nearest neighbor from the set *A_τ,e_*, which we denote *X_τ,e_*(*t*_nn_) for *t* ≠ *t*_nn_. Then, if the distance between these two points in the *e*+1 dimension, *D_τ,e_*_+1_(*t*) = ||*X_τ,e_*_+1_(*t*_nn_)−*X_τ,e_*_+1_(*t*)||_2_, is larger than their distance in the *e* dimension, *D_τ,e_*(*t*) = ||*X_τ,e_*(*t*_nn_) − *X_τ,e_*(*t*)||_2_, these two points are deemed to be false nearest neighbors; specifically, we used a threshold *R* for the ratio between two distances *D_τ,e_*_+1_(*t*)*/D_τ,e_*(*t*) to determine false nearest neighbors. The first embedding dimension *e* that does not have any false nearest neighbors corresponds to the embedding dimension *M* for a given pathogen-country pair. For the main analysis, we used *R* = 10, which was chosen from a sensitivity analysis against simulated data (Supplementary Text). Once we determined the embedding lag *τ* and dimension *M*, we apply the embedding to the entire time series and calculate the nearest neighbor distance against the attractor *A_τ,M_* to obtain a time series of distance from the attractor *D_τ,M_* (*t*).

From a time series of distances from the attractor, we estimated pathogen resilience by fitting a linear regression to an appropriate window. To automatically select fitting windows, we began by smoothing the distance time series using locally estimated scatterplot smoothing (LOESS) to obtain 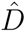*_τ,M_*(*t*), where the smoothing is performed on a log scale and exponentiated afterwards. This smoothing allowed us to find appropriate threshold values for selecting fitting windows that are insensitive to errors in our estimates of distance from the attractor. Then, we determined threshold values (*T*_start_ and *T*_end_) for the smoothed distances and choose the fitting window based on when 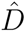*_τ,M_*(*t*) crosses these threshold values for the first time.

These thresholds were determined by first calculating the maximum distance,

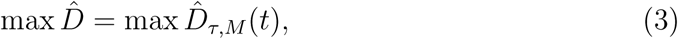

and the mean pre-pandemic distance,

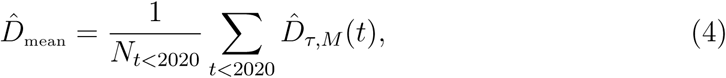

as a reference, and then dividing their ratios into *K* equal bins,

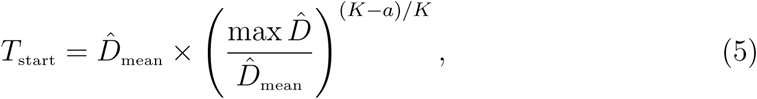

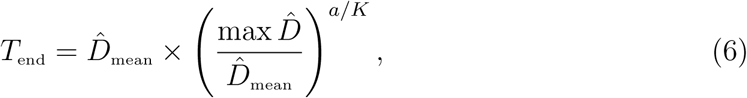

where *a* represents the truncation threshold. This allows us to discard the initial period during which the distance increases (from the introduction of intervention measures) and the final period during which the distance plateaus (as the system reaches an attractor). The fitting window is determined based on when the smoothed distance 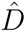*_τ,M_*(*t*) crosses these threshold values for the first time; then, we fit a linear regression to logged (unsmoothed) distances log *D_τ,M_* (*t*) using that window. Alongside the threshold *R* for the false nearest neighbors approach, we tested optimal choices for *K* and *a* values using simulations (Supplementary Text). We used *K* = 19 and *a* = 2 throughout the paper based on the simulation results. We excluded data sets from the regression analysis if the logged average of the estimated distance from the attractor during the last year is at least 2-fold greater than the logged average of the pre-pandemic distance from the attractor.

To evaluate the ability to predict the precise return time for out-of-sample observations, we applied the same framework using data up to the 26th week of 2023. In doing so, we compared the predicted return time with the observed return time, which was determined based on when the smoothed empirical distance from the attractor became close enough to pre-pandemic levels. As a pathogen returns to its pre-pandemic cycle, the the smoothed empirical distance from the attractor is expected to converge to the pre-pandemic levels 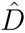_mean_, but the empirical distance may not become smaller than 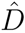_mean_. Therefore, we used 1.2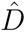_mean_as our thresh-old to obtain an approximate return time. We note that this approach necessarily underestimates the observed return time and therefore should be interpreted as a conservative estimate. For additional validation, we compared these out-of-sample predictions with return time predictions based on full time series.

### Mathematical modeling

Throughout the paper, we use a series of mathematical models to illustrate the concept of pathogen resilience and to understand the determinants of pathogen resilience. In general, the intrinsic resilience of a given system is given by the largest real part of the eigenvalues of the linearized system at endemic equilibrium. Here, we focus on the SIRS model with demography (birth and death) and present the details of other models in Supplementary Text. The SIRS (Susceptible-Infected-Recovered-Susceptible) model is the simplest model that allows for waning of immunity, where recovered (immune) individuals are assumed to become fully susceptible after an average of 1*/δ* time period. The dynamics of the SIRS model is described by the following set of differential equations:

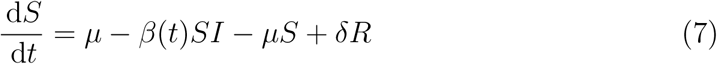

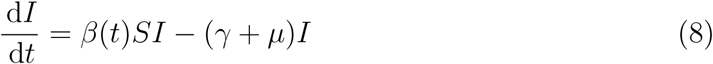

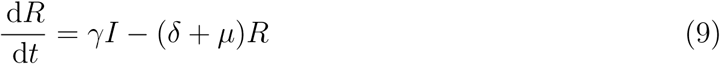

where *µ* represents the birth and death rates, *β*(*t*) represents the time-varying transmission rate, and *γ* represents the recovery rate. The basic reproduction number R_0_(*t*) = *β*(*t*)/(*γ* + *µ*) is defined as the average number of secondary infections that a single infected individual would cause in a fully susceptible population at time *t* and measures the intrinsic transmissibility of a pathogen.

When we introduced the idea of pathogen resilience (Figure 2), we imposed sinusoidal changes to the transmission rate to account for seasonal transmission,

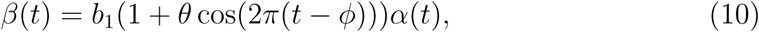

where *b*_1_ represents the baseline transmission rate, *θ* represents the seasonal amplitude, and *ϕ* represents the seasonal offset term. Here, we also introduced an extra multiplicative term *α*(*t*) to account for the impact of pandemic perturbations, where *α*(*t*) *<* 1 indicates transmission reduction. Figure 2A and 2B were generated assuming *b*_1_ = 3 × (365/7 + 1/50)/years, *θ* = 0.2, *ϕ* = 0, *µ* = 1/50/years, *γ* = 365/7/years, and *δ* = 1/2/years. Specifically, *b*_1_ = 3 ×(365/7 + 1/50)/years implies R_0_ = 3, where (365/7 + 1/50)/years represent the rate of recovery. In Figure 2A, we imposed a 50% transmission reduction for 6 months from 2020:

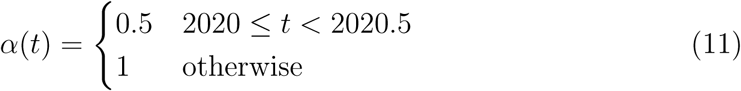

In Figure 2B, we imposed a 50% transmission reduction for 6 months from 2020 and a permanent 10% reduction onward:

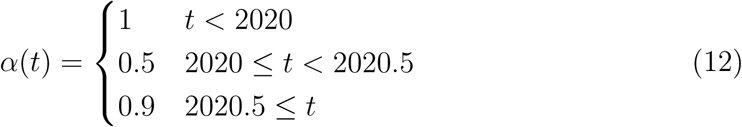

In both scenarios, we simulated the SIRS model from the same initial conditions (*S*(0) = 1/R_0_, *I*(0) = 1 × 10^−6^, and *R*(0) = 1 − *S*(0) − *I*(0)) from 1900 until 2030.

When we include importations (Supplementary Figures S2 and S3), we model the force of infection as *β*(*t*)(*I* + *ω*) instead and vary *ω* between 10^−7^ and 10^−4^ per day. Throughout the paper, all deterministic models were solved using the lsoda solver from the deSolve package [48] in R [49].

To measure the empirical resilience of the SIRS model (Figure 2C and 2F), we computed the normalized distance between post-intervention susceptible and logged infected proportions and their corresponding unperturbed values at the same time:

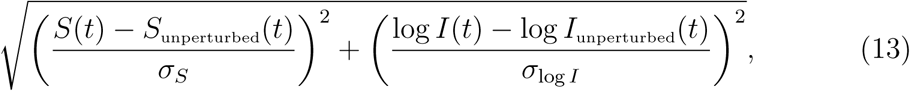

where *σ_S_* and *σ*_log_ *_I_* represent the standard deviation in the unperturbed susceptible and logged infected proportions. The unperturbed values were obtained by simulating the same SIRS model without pandemic perturbations (*α* = 1). We normalized the differences in susceptible and logged infected proportions to allow both quantities to equally contribute to the changes in distance from the attractor. We used logged prevalence, instead of absolute prevalence, in order to capture epidemic dynamics in deep troughs during the intervention period. In Supplementary Materials, we also compared how the degree of seasonal transmission affects empirical resilience by varying *θ* from 0 to 0.4; when we assumed no seasonality (*θ* = 0), we did not normalize the distance because the standard deviation of pre-intervention dynamics are zero.

We used the SIRS model to understand how underlying epidemiological parameters affect pathogen resilience and determine the relationship to underlying susceptible host dynamics. For the simple SIRS model without seasonal transmission (*θ* = 0), the intrinsic resilience equals

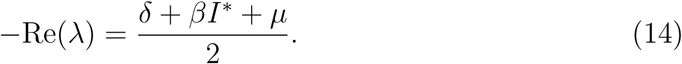

Here, *I*^∗^ represents the prevalence at endemic equilibrium:

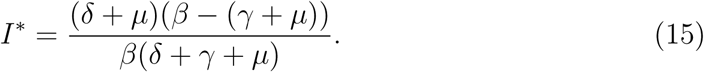

The susceptible replenishment rate is given by

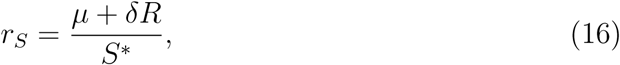

where *S*^∗^ = 1/R_0_ represents the equilibrium proportion of susceptible individuals. We varied the basic reproduction number R_0_ between 1.1 to 6 and the average duration of immunity 1*/δ* between 2 to 4 years, and computed these two quantities. In doing so, we fixed all other parameters: *µ* = 1/80/years and *γ* = 365/7/years. When infection provides a life-long immunity (*δ* = 0), the model collapses to the SIR model and the intrinsic resilience is exactly proportional to the susceptible replenishment rate:

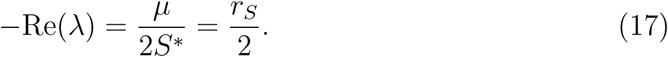

Finally, we used a seasonally unforced stochastic SIRS model without demography to understand how pathogen resilience affects sensitivity of the system to demographic stochasticity (see Supplementary Text for the details of the stochastic SIRS model). By varying the basic reproduction number R_0_ between 2 to 20 and the average duration of immunity 1*/δ* between 1 to 40 years, we ran the SIRS model for 100 years and computed the epidemic amplitude, which we defined as (max *I* − min *I*)/(2*Ī*). Each simulation began from the equilibrium, and we truncated the initial 25 years before computing the epidemic amplitude. In doing so, we assumed *γ* = 365/7/years and fixed the population size to 1 billion to prevent any fadeouts. We also considered a seasonally forced stochastic SIRS model without demography, assuming an amplitude of seasonal forcing of 0.04; in this case, we computed the relative epidemic amplitude by comparing the deterministic and stochastic trajectories (Supplementary Materials).

## Acknowledgement

We thank Anthony R. Ives for helpful discussion. We thank Centers for Disease Control and Prevention, National Respiratory and Enteric Virus Surveillance System (NREVSS) for providing time series data for respiratory infection cases in the US.

## Data availability

All data and code are stored in a publicly available GitHub repository (https://github.com/cobeylab/return-time).

## References

[1] Rachel E Baker, Sang Woo Park, Wenchang Yang, Gabriel A Vecchi, C Jessica E Metcalf, and Bryan T Grenfell. The impact of COVID-19 nonpharmaceutical interventions on the future dynamics of endemic infections. Proceedings of the National Academy of Sciences, 117(48):30547–30553, 2020.

[2] Gabriela B Gomez, Cedric Mahe, and Sandra S Chaves. Uncertain effects of the pandemic on respiratory viruses. Science, 372(6546):1043–1044, 2021.

[3] Mihaly Koltai, Fabienne Krauer, David Hodgson, Edwin van Leeuwen, Marina Treskova-Schwarzbach, Mark Jit, and Stefan Flasche. Determinants of RSV epidemiology following suppression through pandemic contact restrictions. Epidemics, 40:100614, 2022.

[4] Sang Woo Park, Brooklyn Noble, Emily Howerton, Bjarke F Nielsen, Sarah Lentz, Lilliam Ambroggio, Samuel Dominguez, Kevin Messacar, and Bryan T Grenfell. Predicting the impact of non-pharmaceutical interventions against COVID-19 on Mycoplasma pneumoniae in the United States. Epidemics, 49:100808, 2024.

[5] Amanda C Perofsky, Chelsea L Hansen, Roy Burstein, Shanda Boyle, Robin Prentice, Cooper Marshall, David Reinhart, Ben Capodanno, Melissa Truong, Kristen Schwabe-Fry, et al. Impacts of human mobility on the citywide transmission dynamics of 18 respiratory viruses in pre-and post-COVID-19 pandemic years. Nature communications, 15(1):4164, 2024.

[6] Eric J Chow, Timothy M Uyeki, and Helen Y Chu. The effects of the COVID-19 pandemic on community respiratory virus activity. Nature Reviews Microbiology, 21(3):195–210, 2023.

[7] Stephen M Kissler, Christine Tedijanto, Edward Goldstein, Yonatan H Grad, and Marc Lipsitch. Projecting the transmission dynamics of SARS-CoV-2 through the postpandemic period. Science, 368(6493):860–868, 2020.

[8] Rachel E Baker, Chadi M Saad-Roy, Sang Woo Park, Jeremy Farrar, C Jessica E Metcalf, and Bryan T Grenfell. Long-term benefits of nonpharmaceutical interventions for endemic infections are shaped by respiratory pathogen dynamics. Proceedings of the National Academy of Sciences, 119(49):e2208895119, 2022.

[9] Maaike C Swets, Clark D Russell, Ewen M Harrison, Annemarie B Docherty, Nazir Lone, Michelle Girvan, Hayley E Hardwick, Leonardus G Visser, Peter JM Openshaw, Geert H Groeneveld, et al. SARS-CoV-2 co-infection with influenza viruses, respiratory syncytial virus, or adenoviruses. The Lancet, 399(10334):1463–1464, 2022.

[10] Chun-Yang Lin, Joshua Wolf, David C Brice, Yilun Sun, Macauley Locke, Sean Cherry, Ashley H Castellaw, Marie Wehenkel, Jeremy Chase Crawford, Veronika I Zarnitsyna, et al. Pre-existing humoral immunity to human common cold coronaviruses negatively impacts the protective SARS-CoV-2 antibody response. Cell host & microbe, 30(1):83–96, 2022.

[11] Sam M Murray, Azim M Ansari, John Frater, Paul Klenerman, Susanna Dunachie, Eleanor Barnes, and Ane Ogbe. The impact of pre-existing crossreactive immunity on SARS-CoV-2 infection and vaccine responses. Nature Reviews Immunology, 23(5):304–316, 2023.

[12] John-Sebastian Eden, Chisha Sikazwe, Ruopeng Xie, Yi-Mo Deng, Sheena G Sullivan, Alice Michie, Avram Levy, Elena Cutmore, Christopher C Blyth, Philip N Britton, et al. Off-season RSV epidemics in Australia after easing of COVID-19 restrictions. Nature communications, 13(1):2884, 2022.

[13] Stuart L Pimm. The structure of food webs. Theoretical population biology, 16(2):144–158, 1979.

[14] Michael G Neubert and Hal Caswell. Alternatives to resilience for measuring the responses of ecological systems to perturbations. Ecology, 78(3):653–665, 1997.

[15] Lance H Gunderson. Ecological resilience—in theory and application. Annual review of ecology and systematics, 31(1):425–439, 2000.

[16] Vasilis Dakos and Sonia Kefi. Ecological resilience: what to measure and how. Environmental Research Letters, 17(4):043003, 2022.

[17] Floris Takens. Detecting strange attractors in turbulence. In Dynamical Systems and Turbulence, Warwick 1980: proceedings of a symposium held at the University of Warwick 1979/80, pages 366–381. Springer, 2006.

[18] Jonathan Dushoff, Joshua B Plotkin, Simon A Levin, and David JD Earn. Dynamical resonance can account for seasonality of influenza epidemics. Proceedings of the National Academy of Sciences, 101(48):16915–16916, 2004.

[19] Alan Hastings, Carole L Hom, Stephen Ellner, Peter Turchin, and H Charles J Godfray. Chaos in ecology: is mother nature a strange attractor? Annual review of ecology and systematics, pages 1–33, 1993.

[20] Alan Hastings, Karen C Abbott, Kim Cuddington, Tessa Francis, Gabriel Gellner, Ying-Cheng Lai, Andrew Morozov, Sergei Petrovskii, Katherine Scranton, and Mary Lou Zeeman. Transient phenomena in ecology. Science, 361(6406):eaat6412, 2018.

[21] Matthew B Kennel, Reggie Brown, and Henry DI Abarbanel. Determining embedding dimension for phase-space reconstruction using a geometrical construction. Physical review A, 45(6):3403, 1992.

[22] Eugene Tan, Shannon Algar, Débora Corrêa, Michael Small, Thomas Stemler, and David Walker. Selecting embedding delays: An overview of embedding techniques and a new method using persistent homology. Chaos: An Interdisciplinary Journal of Nonlinear Science, 33(3), 2023.

[23] Heungwon Park, William Pontius, Calin C Guet, John F Marko, Thierry Emonet, and Philippe Cluzel. Interdependence of behavioural variability and response to small stimuli in bacteria. Nature, 468(7325):819–823, 2010.

[24] Samantha Bosis, Susanna Esposito, HGM Niesters, GV Zuccotti, G Marseglia, Marcello Lanari, Giovanna Zuin, C Pelucchi, ADME Osterhaus, and Nicola Principi. Role of respiratory pathogens in infants hospitalized for a first episode of wheezing and their impact on recurrences. Clinical microbiology and infection, 14(7):677–684, 2008.

[25] Miguel L O’Ryan, Yalda Lucero, Valeria Prado, María Elena Santolaya, Marcela Rabello, Yanahara Solis, Daniela Berríos, Miguel A O’Ryan-Soriano, Hector Cortés, and Nora Mamani. Symptomatic and asymptomatic rotavirus and norovirus infections during infancy in a Chilean birth cohort. The Pediatric infectious disease journal, 28(10):879–884, 2009.

[26] Virginia E Pitzer, Cécile Viboud, Wladimir J Alonso, Tanya Wilcox, C Jessica Metcalf, Claudia A Steiner, Amber K Haynes, and Bryan T Grenfell. Environmental drivers of the spatiotemporal dynamics of respiratory syncytial virus in the United States. PLoS pathogens, 11(1):e1004591, 2015.

[27] Arthur WD Edridge, Joanna Kaczorowska, Alexis CR Hoste, Margreet Bakker, Michelle Klein, Katherine Loens, Maarten F Jebbink, Amy Matser, Cormac M Kinsella, Paloma Rueda, et al. Seasonal coronavirus protective immunity is short-lasting. Nature medicine, 26(11):1691–1693, 2020.

[28] Jennifer M Radin, Anthony W Hawksworth, Peter E Kammerer, Melinda Balansay, Rema Raman, Suzanne P Lindsay, and Gary T Brice. Epidemiology of pathogen-specific respiratory infections among three US populations. PLoS One, 9(12):e114871, 2014.

[29] Guojian Lv, Limei Shi, Yi Liu, Xuecheng Sun, and Kai Mu. Epidemiological characteristics of common respiratory pathogens in children. Scientific Reports, 14(1):16299, 2024.

[30] Saverio Caini, Adam Meijer, Marta C Nunes, Laetitia Henaff, Malaika Zounon, Bronke Boudewijns, Marco Del Riccio, and John Paget. Probable extinction of influenza b/yamagata and its public health implications: a systematic literature review and assessment of global surveillance databases. The Lancet Microbe, 2024.

[31] Zhiyuan Chen, Joseph L-H Tsui, Bernardo Gutierrez, Simon Busch Moreno, Louis du Plessis, Xiaowei Deng, Jun Cai, Sumali Bajaj, Marc A Suchard, Oliver G Pybus, et al. COVID-19 pandemic interventions reshaped the global dispersal of seasonal influenza viruses. Science, 386(6722):eadq3003, 2024.

[32] Bjarke Frost Nielsen, Sang Woo Park, Emily Howerton, Olivia Frost Lorentzen, Mogens H Jensen, and Bryan T Grenfell. Complex multiannual cycles of Mycoplasma pneumoniae: persistence and the role of stochasticity. arXiv, 2025.

[33] Hai Nguyen-Tran, Sang Woo Park, Kevin Messacar, Samuel R Dominguez, Matthew R Vogt, Sallie Permar, Perdita Permaul, Michelle Hernandez, Daniel C Douek, Adrian B McDermott, et al. Enterovirus D68: a test case for the use of immunological surveillance to develop tools to mitigate the pandemic potential of emerging pathogens. The Lancet Microbe, 3(2):e83–e85, 2022.

[34] Alun L Lloyd. Realistic distributions of infectious periods in epidemic models: changing patterns of persistence and dynamics. Theoretical population biology, 60(1):59–71, 2001.

[35] Jennie S Lavine, Aaron A King, and Ottar N Bjørnstad. Natural immune boosting in pertussis dynamics and the potential for long-term vaccine failure. Proceedings of the National Academy of Sciences, 108(17):7259–7264, 2011.

[36] Nicholas G Reich, Sourya Shrestha, Aaron A King, Pejman Rohani, Justin Lessler, Siripen Kalayanarooj, In-Kyu Yoon, Robert V Gibbons, Donald S Burke, and Derek AT Cummings. Interactions between serotypes of dengue highlight epidemiological impact of cross-immunity. Journal of The Royal Society Interface, 10(86):20130414, 2013.

[37] Chadi M Saad-Roy, Simon A Levin, Bryan T Grenfell, and Mike Boots. Epidemiological impacts of post-infection mortality. Proceedings of the Royal Society B, 290(2002):20230343, 2023.

[38] Tim G Benton and Alastair Grant. Elasticity analysis as an important tool in evolutionary and population ecology. Trends in ecology & evolution, 14(12):467– 471, 1999.

[39] Amy Matser, Nienke Hartemink, Hans Heesterbeek, Alison Galvani, and Stephen Davis. Elasticity analysis in epidemiology: an application to tick-borne infections. Ecology Letters, 12(12):1298–1305, 2009.

[40] Sarah Cobey and Edward B Baskerville. Limits to causal inference with statespace reconstruction for infectious disease. PloS one, 11(12):e0169050, 2016.

[41] P Rohani, CJ Green, NB Mantilla-Beniers, and Bryan T Grenfell. Ecological interference between fatal diseases. Nature, 422(6934):885–888, 2003.

[42] Sema Nickbakhsh, Colette Mair, Louise Matthews, Richard Reeve, Paul CD Johnson, Fiona Thorburn, Beatrix Von Wissmann, Arlene Reynolds, James McMenamin, Rory N Gunson, et al. Virus–virus interactions impact the population dynamics of influenza and the common cold. Proceedings of the National Academy of Sciences, 116(52):27142–27150, 2019.

[43] Public Health Agency of Canada. Respiratory virus detections in Canada. 2024. https://www.canada.ca/en/public-health/services/surveillance/respiratory-virus-detections-canada.html. Accessed Sep 4, 2024.

[44] Centre for Health Protection. Detection of pathogens from respiratory specimens. 2024. https://www.chp.gov.hk/en/statistics/data/10/641/642/2274.html. Accessed Nov 20, 2024.

[45] Centre for Health Protection. Detection of gastroenteritis viruses from faecal specimens. 2024. https://www.chp.gov.hk/en/statistics/data/10/641/717/3957.html. Accessed Nov 20, 2024.

[46] Korea Disease Control and Prevention Agency. Acute respiratory infection, Sentinel surveillance infectious diseases. 2024. https://dportal.kdca.go.kr/pot/is/st/ari.do. Accessed Nov 20, 2024.

[47] Edward Goldstein, Sarah Cobey, Saki Takahashi, Joel C Miller, and Marc Lipsitch. Predicting the epidemic sizes of influenza A/H1N1, A/H3N2, and B: a statistical method. PLoS medicine, 8(7):e1001051, 2011.

[48] Karline Soetaert, Thomas Petzoldt, and R Woodrow Setzer. Solving differential equations in R: package deSolve. Journal of statistical software, 33:1–25, 2010.

[49] R Core Team. R: A Language and Environment for Statistical Computing. R Foundation for Statistical Computing, Vienna, Austria, 2023.

