## supplementary file for "Susceptible host dynamics explain pathogen resilience to perturbations"

### Supplementary Text

#### Resilience of a stage-structured system.

In Figure 2G–I, we used a more realistic, stage-structured model to illustrate how transient phenomena can cause the system to slow down. Specifically, we used the stage-structured RSV model proposed by [1], which assumes that subsequent reinfections cause an individual to become less susceptible and transmissible than previous infections. In contrast to a standard SIRS model, this model does not include a recovered compartment, which allow for temporary protection against new infections, and assumes that recoverd individuals are immediately susceptible to new infections. The model dynamics can be described as follows:

$$\frac{\mathrm{d}M}{\mathrm{d}t} = \mu - (\omega + \mu)M \tag{S1}$$

$$\frac{\mathrm{d}S_0}{\mathrm{d}t} = \omega M - (\lambda(t) + \mu)S_0 \tag{S2}$$

$$\frac{\mathrm{d}I_1}{\mathrm{d}t} = \lambda(t)S_0 - (\gamma_1 + \mu)I_1 \tag{S3}$$

$$\frac{\mathrm{d}S_1}{\mathrm{d}t} = \gamma_1 I_1 - (\sigma_1\lambda(t) + \mu)S_1 \tag{S4}$$

$$\frac{\mathrm{d}I_2}{\mathrm{d}t} = \sigma_1\lambda(t)S_1 - (\gamma_2 + \mu)I_2 \tag{S5}$$

$$\frac{\mathrm{d}S_2}{\mathrm{d}t} = \gamma_2 I_2 - (\sigma_2\lambda(t) + \mu)S_2 \tag{S6}$$

$$\frac{\mathrm{d}I_3}{\mathrm{d}t} = \sigma_2\lambda(t)S_2 - (\gamma_3 + \mu)I_3 \tag{S7}$$

$$\frac{\mathrm{d}S_3}{\mathrm{d}t} = \gamma_3 I_3 - (\sigma_3\lambda(t) + \mu)S_3 + \gamma_4 I_4 \tag{S8}$$

$$\frac{\mathrm{d}I_4}{\mathrm{d}t} = \sigma_3\lambda(t)S_3 - (\gamma_4 + \mu)I_4 \tag{S9}$$

where $M$ represents the proportion of individuals who are maternally immune; $S_i$ represents the proportion of individuals who are susceptible after $i$ prior infections; $I_i$ represents the proportion of individuals who are currently (re)-infected with their $i$-th infection; $\mu$ represents the birth and death rates; $1/\omega$ represents the mean duration of maternal immunity; $1/\gamma_i$ represents the mean duration of infection; $\lambda(t)$ represents the force of infection; and $\sigma_i$ represents the reduction in susceptibility for the $i$-th reinfection. The force of infection is modeled using a sinusoidal function:

$$\beta(t) = b_1(1 + \theta\cos(2\pi(t - \phi)))\alpha(t) \tag{S10}$$

$$\lambda(t) = \beta(I_1 + \rho_1 I_2 + \rho_2(I_3 + I_4)), \tag{S11}$$

where $b_1$ represents the baseline transmission rate; $\theta$ represents the seasonal amplitude; $\phi$ represents the seasonal offset term; $\alpha(t)$ represents the intervention effect;

and $\rho_i$ represents the impact of immunity on transmission reduction. We used the following parameters to simulate the impact of interventions on epidemic dynamics [1]: $b_1 = 9 \times (365/10 + 1/80)$/years, $\theta = 0.2$, $\phi = -0.1$, $\omega = 365/112$/years, $\gamma_1 = 365/10$/years, $\gamma_2 = 365/7$/years, $\gamma_3 = 365/5$/years, $\sigma_1 = 0.76$, $\sigma_2 = 0.6$, $\sigma_3 = 0.4$, $\rho_1 = 0.75$, $\rho_2 = 0.51$, and $\mu = 1/80$/years. We assumed a 50% transmission reduction for 6 months from 2020:

$$\alpha(t) = \begin{cases} 0.5 & 2020 \leq t < 2020.5 \\ 1 & \text{otherwise} \end{cases} \tag{S12}$$

The model was simulated from 1900 to 2030 using the following initial conditions: $M = 0$, $S_0 = 1/\mathcal{R}_0 - I_1$, $I_1 = 1 \times 10^{-6}$, $S_1 = 1 - 1/\mathcal{R}_0$, $I_2 = 0$, $S_2 = 0$, $I_3 = 0$, $S_3 = 0$, and $I_4 = 0$. For the phase plane analysis (Figure 2H) and distance analysis (Figure 2I), we relied on the average susceptibility,

$$\bar{S} = S_0 + \sigma_1 S_1 + \sigma_2 S_2 + \sigma_3 S_3, \tag{S13}$$

and total prevalence,

$$I_{\text{total}} = I_1 + I_2 + I_3 + I_4. \tag{S14}$$

These quantities were used to compute the normalized distance from the attractor, as described in the main text.

We note that this system, without seasonally forced transmission rates, has 9 eigenvalues: -73.01, -53.11, -38.90, -3.27, -0.82+2.57i, -0.82-2.57i, -1.61, -1.18, and -0.01 (in the unit of 1/years). While the eigenvalue -0.01 has the largest real part, the magnitude is too close to 0 for the impact of this eigenvalue to be reflected in our resilience estimates. Instead, we chose real parts of the eigenvalues $-0.82 \pm 2.57i$ as our intrinsic resilience for this system and plotted them in Figure 2I; as we can see in this figure, this value captures the return rate of the system to the attractor.

### Resilience of a multistrain system.

We used a simple two-strain model to show that a multistrain host-pathogen system that is coupled through cross immunity can be described by a single resilience value. The model dynamics can be described as follows [2]:

$$\frac{\mathrm{d}S}{\mathrm{d}t} = \mu - \mu S - (\lambda_1 + \lambda_2) S + \delta_1 R_1 + \delta_2 R_2 \tag{S15}$$

$$\frac{\mathrm{d}I_1}{\mathrm{d}t} = \lambda_1 S - (\gamma_1 + \mu) I_1 \tag{S16}$$

$$\frac{\mathrm{d}I_2}{\mathrm{d}t} = \lambda_2 S - (\gamma_2 + \mu) I_2 \tag{S17}$$

$$\frac{\mathrm{d}R_1}{\mathrm{d}t} = \gamma_1 I_1 - \lambda_2 \epsilon_{21} R_1 - \delta_1 R_1 + \delta_2 R - \mu R_1 \tag{S18}$$

$$\frac{\mathrm{d}R_2}{\mathrm{d}t} = \gamma_2 I_2 - \lambda_1 \epsilon_{12} R_2 - \delta_2 R_2 + \delta_1 R - \mu R_2 \tag{S19}$$

$$\frac{\mathrm{d}J_1}{\mathrm{d}t} = \lambda_1 \epsilon_{12} R_2 - (\gamma_1 + \mu) J_1 \tag{S20}$$

$$\frac{\mathrm{d}J_2}{\mathrm{d}t} = \lambda_2 \epsilon_{21} R_1 - (\gamma_2 + \mu) J_2 \tag{S21}$$

$$\frac{\mathrm{d}R}{\mathrm{d}t} = \gamma_1 J_1 + \gamma_2 J_2 - \delta_1 R - \delta_2 R - \mu R \tag{S22}$$

where $S$ represents the proportion of individuals who are fully susceptible to infections by both strains; $I_1$ represents the proportion of individuals who are infected with strain 1 without prior immunity; $I_2$ represents the proportion of individuals who are infected with strain 2 without prior immunity; $R_1$ represents the proportion of individuals who are fully immune against strain 1 and partially susceptible to reinfection by strain 2; $R_2$ represents the proportion of individuals who are fully immune against strain 2 and partially susceptible to reinfection by strain 1; $J_1$ represents the proportion of individuals who are infected with strain 1 with prior immunity against strain 2; $J_2$ represents the proportion of individuals who are infected with strain 2 with prior immunity against strain 1; $R$ represents the proportion of individuals who are immune to infections from both strains; $\mu$ represents the birth/death rate; $\lambda_1$ and $\lambda_2$ represent the force of infection from strains 1 and 2, respectively; $\delta_1$ and $\delta_2$ represent the waning immunity rate; $\gamma_1$ and $\gamma_2$ represent the recovery rate; $\epsilon_{21}$ and $\epsilon_{12}$ represent the susceptibility to reinfection with strains 2 and 1, respectively, given prior immunity from infection with strains 1 and 2, respectively. The force of infection is modeled as follows:

$$\beta_1 = b_1(1 + \theta_1 \sin(2\pi(t - \phi_1)))\alpha(t) \tag{S23}$$

$$\beta_2 = b_2(1 + \theta_2 \sin(2\pi(t - \phi_2)))\alpha(t) \tag{S24}$$

$$\lambda_1 = \beta_1(I_1 + J_1) \tag{S25}$$

$$\lambda_2 = \beta_2(I_2 + J_2) \tag{S26}$$

In Supplementary Figures S2–S4, we assumed the following parameters: $b_1 = 2 \times 52$/years, $b_2 = 4 \times 52$/years, $\phi_1 = \phi_2 = 0$, $\epsilon_{12} = 0.9$, $\epsilon_{21} = 0.5$, $\gamma_1 = \gamma_2 = 52$/years, $\delta_1 = \delta_2 = 1$/years, and $\mu = 1/70$/years. For all simulations, we assumed a 50% transmission reduction for 6 months from 2020:

$$\alpha(t) = \begin{cases} 0.5 & 2020 \leq t < 2020.5 \\ 1 & \text{otherwise} \end{cases} \tag{S27}$$

The seasonal amplitude $\theta$ is varied from 0 to 0.4. All simulations were run from 1900 to 2030 with following initial conditions: $S(0) = 1 - 2 \times 10^{-6}$, $I_1(0) = 1 \times 10^{-6}$, $I_2(0) = 1 \times 10^{-6}$, $R_1(0) = 0$, $R_2(0) = 0$, $J_1(0) = 0$, $J_2(0) = 0$, and $R(0) = 0$.

We considered three scenarios for measuring pathogen resilience: (1) we only have information about strain 1, (2) we only have information about strain 2, and (3) we are unable to distinguish between strains. In the first two scenarios (see panels A–C for strain 1 and panels D–F for strain 2), we considered the dynamics of average susceptibility for each strain and total prevalence:

$$\bar{S}_1 = S + \epsilon_{12} R_2 \tag{S28}$$

$$\hat{I}_1 = I_1 + J_1 \tag{S29}$$

$$\bar{S}_2 = S + \epsilon_{21} R_1 \tag{S30}$$

$$\hat{I}_2 = I_2 + J_2 \tag{S31}$$

In the third scenario (panels G–I), we considered the dynamics of total susceptible and infected populations:

$$\hat{S} = S + R_2 + R_1 \tag{S32}$$

$$\hat{I} = I_1 + J_1 + I_2 + J_2 \tag{S33}$$

These quantities were used to compute the normalized distance from the attractor, as described in the main text.

#### Estimating intrinsic resilience using a mechanistic model

We tested whether we can reliably estimate the intrinsic resilience of a system by fitting a mechanistic model. Specifically, we simulated case time series from stochastic SIRS and two-strain models and fitted a simple, deterministic SIRS model using a Bayesian framework [3, 4, 5].

We simulated the models in discrete time with a daily time step ($\Delta t = 1$), incorporating demographic stochasticity:

$$\beta(t) = \mathcal{R}_0 \left(1 + \theta \cos\left(\frac{2\pi t}{364}\right)\right) \alpha(t)(1 - \exp(-\gamma)) \tag{S34}$$

$$\mathrm{FOI}(t) = \beta(t) I(t - \Delta t)/N \tag{S35}$$

$$B(t) \sim \mathrm{Poisson}(\mu N) \tag{S36}$$

$$\Delta S(t) \sim \mathrm{Binom}\left(S(t - \Delta t), 1 - \exp(-(\mathrm{FOI}(t) + \mu)\Delta t)\right) \tag{S37}$$

$$N_{SI}(t) \sim \mathrm{Binom}\left(\Delta S(t), \frac{\mathrm{FOI}(t)}{\mathrm{FOI}(t) + \mu}\right) \tag{S38}$$

$$\Delta I(t) \sim \mathrm{Binom}\left(I(t - \Delta t), 1 - \exp(-(\gamma + \mu)\Delta t)\right) \tag{S39}$$

$$N_{IR}(t) \sim \mathrm{Binom}\left(\Delta I(t), \frac{\gamma}{\gamma + \mu}\right) \tag{S40}$$

$$\Delta R(t) \sim \mathrm{Binom}\left(R(t - \Delta t), 1 - \exp(-(\delta + \mu)\Delta t)\right) \tag{S41}$$

$$N_{RS}(t) \sim \mathrm{Binom}\left(\Delta R(t), \frac{\delta}{\delta + \mu}\right) \tag{S42}$$

$$S(t) = S(t - \Delta t) + N_{RS}(t) + B(t) - \Delta S(t) \tag{S43}$$

$$I(t) = I(t - \Delta t) + N_{SI}(t) - \Delta I(t) \tag{S44}$$

$$R(t) = R(t - \Delta t) + N_{IR}(t) - \Delta R(t) \tag{S45}$$

where FOI represents the force of infection; $N_{ij}$ represents the number of individuals moving from compartment $i$ to $j$ on a given day; and $B(t)$ represents the number of new births. All other parameters definitions can be found in the description of the deterministic version of the model. We simulated the model on a daily scale—assuming 364 days in a year so that it can be evenly grouped into 52 weeks—with the following parameters: $\mathcal{R}_0 = 3$, $\theta = 0.1$, $\gamma = 1/7$/days, $\delta = 1/(364 \times 2)$/days, $\mu = 1/(364 \times 50)$/days, and $N = 1 \times 10^8$. The model was simulated from 1900 to 2030 assuming $S(0) = N/3$, $I(0) = 100$, and $R(0) = N - S(0) - I(0)$. The observed incidence from the model was then simulated as follows:

$$C(t) = \text{Beta-Binom}(N_{SI}(t), \rho, k), \tag{S46}$$

where $\rho$ represents the reporting probability and $k$ represents the overdispersion parameter of beta-binomial distribution. Here, we used the beta-binomial distribution to account for overdispersion in reporting. We assumed $\rho = 0.002$ (i.e., 0.2% probability) and $k = 1000$.

We used an analogous approach for the two-strain model:

$$\beta_1(t) = b_1 \left(1 + \theta_1 \cos\left(\frac{2\pi(t - \phi_1)}{364}\right)\right) \alpha(t) \tag{S47}$$

$$\beta_2(t) = b_2 \left(1 + \theta_2 \cos\left(\frac{2\pi(t - \phi_2)}{364}\right)\right) \alpha(t) \tag{S48}$$

$$\text{FOI}_1(t) = \beta_1(t)(I_1(t - \Delta t) + J_1(t - \Delta t))/N \tag{S49}$$

$$\text{FOI}_2(t) = \beta_2(t)(I_2(t - \Delta t) + J_2(t - \Delta t))/N \tag{S50}$$

$$B(t) \sim \text{Poisson}(\mu N) \tag{S51}$$

$$\Delta S(t) \sim \text{Binom}\left(S(t - \Delta t), 1 - \exp(-(\text{FOI}_1(t) + \text{FOI}_2(t) + \mu)\Delta t)\right) \tag{S52}$$

$$N_{SI_1}(t) \sim \text{Binom}\left(\Delta S(t), \frac{\text{FOI}_1(t)}{\text{FOI}_1(t) + \text{FOI}_2(t) + \mu}\right) \tag{S53}$$

$$N_{SI_2}(t) \sim \text{Binom}\left(\Delta S(t), \frac{\text{FOI}_2(t)}{\text{FOI}_1(t) + \text{FOI}_2(t) + \mu}\right) \tag{S54}$$

$$\Delta I_1(t) \sim \text{Binom}\left(I_1(t - \Delta t), 1 - \exp(-(\gamma_1 + \mu)\Delta t)\right) \tag{S55}$$

$$N_{I_1R_1}(t) \sim \text{Binom}\left(\Delta I_1(t), \frac{\gamma_1}{\gamma_1 + \mu}\right) \tag{S56}$$

$$\Delta I_2(t) \sim \text{Binom}\left(I_2(t - \Delta t), 1 - \exp(-(\gamma_2 + \mu)\Delta t)\right) \tag{S57}$$

$$N_{I_2R_2}(t) \sim \text{Binom}\left(\Delta I_2(t), \frac{\gamma_2}{\gamma_2 + \mu}\right) \tag{S58}$$

$$\Delta R_1(t) \sim \text{Binom}\left(R_1(t - \Delta t), 1 - \exp(-(\epsilon_{21}\text{FOI}_2(t) + \rho_1 + \mu)\Delta t)\right) \tag{S59}$$

$$N_{R_1S}(t) \sim \text{Binom}\left(\Delta R_1(t), \frac{\rho_1}{\epsilon_{21}\text{FOI}_2(t) + \rho_1 + \mu}\right) \tag{S60}$$

$$N_{R_1J_2}(t) \sim \text{Binom}\left(\Delta R_1(t), \frac{\epsilon_{21}\text{FOI}_2(t)}{\epsilon_{21}\text{FOI}_2(t) + \rho_1 + \mu}\right) \tag{S61}$$

$$\Delta R_2(t) \sim \text{Binom}\left(R_2(t-\Delta t), 1-\exp(-(\epsilon_{12}\text{FOI}_1(t)+\rho_2+\mu)\Delta t)\right) \tag{S62}$$

$$N_{R_2S}(t) \sim \text{Binom}\left(\Delta R_2(t), \frac{\rho_2}{\epsilon_{12}\text{FOI}_1(t)+\rho_2+\mu}\right) \tag{S63}$$

$$N_{R_2J_1}(t) \sim \text{Binom}\left(\Delta R_2(t), \frac{\epsilon_{12}\text{FOI}_1(t)}{\epsilon_{12}\text{FOI}_1(t)+\rho_2+\mu}\right) \tag{S64}$$

$$\Delta J_1(t) \sim \text{Binom}\left(J_1(t-\Delta t), 1-\exp(-(\gamma_1+\mu)\Delta t)\right) \tag{S65}$$

$$N_{J_1R}(t) \sim \text{Binom}\left(\Delta J_1(t), \frac{\gamma_1}{\gamma_1+\mu}\right) \tag{S66}$$

$$\Delta J_2(t) \sim \text{Binom}\left(J_2(t-\Delta t), 1-\exp(-(\gamma_2+\mu)\Delta t)\right) \tag{S67}$$

$$N_{J_2R}(t) \sim \text{Binom}\left(\Delta J_2(t), \frac{\gamma_2}{\gamma_2+\mu}\right) \tag{S68}$$

$$\Delta R(t) \sim \text{Binom}\left(R(t-\Delta t), 1-\exp(-(\rho_1+\rho_2+\mu)\Delta t)\right) \tag{S69}$$

$$N_{RR_1}(t) \sim \text{Binom}\left(\Delta R(t), \frac{\rho_1}{\rho_1+\rho_2+\mu}\right) \tag{S70}$$

$$N_{RR_2}(t) \sim \text{Binom}\left(\Delta R(t), \frac{\rho_2}{\rho_1+\rho_2+\mu}\right) \tag{S71}$$

$$S(t) = S(t-\Delta t) - \Delta S(t) + B(t) + N_{R_1S}(t) + N_{R_2S}(t) \tag{S72}$$

$$I_1(t) = I_1(t-\Delta t) - \Delta I_1(t) + N_{SI_1}(t) \tag{S73}$$

$$I_2(t) = I_2(t-\Delta t) - \Delta I_2(t) + N_{SI_2}(t) \tag{S74}$$

$$R_1(t) = R_1(t-\Delta t) - \Delta R_1(t) + N_{I_1R_1}(t) + N_{RR_1}(t) \tag{S75}$$

$$R_2(t) = R_2(t-\Delta t) - \Delta R_2(t) + N_{I_2R_2}(t) + N_{RR_2}(t) \tag{S76}$$

$$J_1(t) = J_1(t-\Delta t) - \Delta J_1(t) + N_{R_2J_1}(t) \tag{S77}$$

$$J_2(t) = J_2(t-\Delta t) - \Delta J_2(t) + N_{R_1J_2}(t) \tag{S78}$$

$$R(t) = R(t-\Delta t) - \Delta R(t) + N_{J_1R}(t) + N_{J_2R}(t) \tag{S79}$$

We simulated the model on a daily scale with previously estimated parameters for the RSV-HMPV interaction [2]: $b_1 = 1.7$/weeks, $b_1 = 1.95$/weeks, $\theta_1 = 0.4$, $\theta_2 = 0.3$, $\phi_1 = 0.005 \times 7/364$, $\phi_2 = 4.99 \times 7/364$, $\epsilon_{12} = 0.92$, $\epsilon_{21} = 0.45$, $\gamma_1 = 1/10$/days, $\gamma_2 = 1/10$/days, $\rho_1 = 1/364$/days, $\rho_2 = 1/364$/days, $\mu = 1/(70 \times 364)$/days, and $N = 1 \times 10^8$. The model was simulated from 1900 to 2030 assuming $S(0) = N - 200$, $I_1(0) = 100$, $I_2(0) = 100$, $R_1(0) = 0$, $R_2(0) = 0$, $J_1(0) = 0$, $J_2(0) = 0$, and $R(0) = 0$. The observed incidence for each strain is then simulated as follows:

$$C_1(t) = \text{Beta-Binom}(N_{SI_1}(t) + N_{R_2J_1}, \rho, k), \tag{S80}$$

$$C_2(t) = \text{Beta-Binom}(N_{SI_2}(t) + N_{R_1J_2}, \rho, k), \tag{S81}$$

where $\rho$ represents the reporting probability and $k$ represents the overdispersion parameter of beta-binomial distribution. We assumed $\rho = 0.002$ (i.e., 0.2% probability) and $k = 500$. We also considered the total incidence: $C_{\text{total}}(t) = C_1(t) + C_2(t)$.

For both models, we considered a more realistically shaped pandemic perturbation $\alpha(t)$ to challenge our ability to estimate the intervention effects. Thus, we assumed a 40% transmission reduction for 3 months from March 2020, followed by a 10% transmission reduction for 6 months, 20% transmission reduction for 3 months, and a final return to normal levels:

$$\alpha(t) = \begin{cases} 1 & t < 2020.25 \\ 0.6 & 2020.25 \leq t < 2020.5 \\ 0.9 & 2020.5 \leq t < 2021 \\ 0.8 & 2021 \leq t < 2021.25 \\ 1 & 2021.25 \leq t \end{cases}. \tag{S82}$$

For all simulations, we truncated the time series from the beginning of 2014 to the end of 2023 and aggregated them into weekly cases.

To infer intrinsic resilience from time series, we fitted a simple discrete time, deterministic SIRS model in a Bayesian framework [3]:

$$\Delta t = 1\,\text{week} \tag{S83}$$
$$\text{FOI}(t) = \beta(t)(I(t-\Delta t)+\omega)/N \tag{S84}$$
$$\Delta S(t) = [1-\exp(-(\text{FOI}(t)+\mu)\Delta t)]\,S(t-\Delta t) \tag{S85}$$
$$N_{SI}(t) = \frac{\text{FOI}(t)\Delta S(t)}{\text{FOI}(t)+\mu} \tag{S86}$$
$$\Delta I(t) = [1-\exp(-(\gamma+\mu)\Delta t)]\,I(t-\Delta t) \tag{S87}$$
$$N_{IR}(t) = \frac{\gamma \Delta I(t)}{\gamma+\mu} \tag{S88}$$
$$\Delta R(t) = [1-\exp(-(\nu+\mu)\Delta t)]\,R(t-\Delta t) \tag{S89}$$
$$N_{RS}(t) = \frac{\nu \Delta R(t)}{\nu+\mu} \tag{S90}$$
$$S(t) = S(t-\Delta t) + \mu S - \Delta S(t) + N_{RS}(t) \tag{S91}$$
$$I(t) = I(t-\Delta t) - \Delta I(t) + N_{SI}(t) \tag{S92}$$
$$R(t) = R(t-\Delta t) - \Delta R(t) + N_{IR}(t) \tag{S93}$$

where we include an extra term $\omega$ to account for importation. Although actual simulations did not include any importation, we had found that including this term generally helped with model convergence in previous analyses [3]. The transmission rate was divided into a seasonal term $\beta_{\text{seas}}(t)$ (repeated every year) and intervention term $\alpha(t)$, which were estimated jointly:

$$\beta(t) = \beta_{\text{seas}}(t)\alpha(t), \tag{S94}$$

where $\alpha < 1$ corresponds to reduction in transmission due to intervention effects. To constrain the smoothness of $\beta_{\text{seas}}(t)$, we imposed cyclic, random-walk priors:

$$\beta_{\text{seas}}(t) \sim \text{Normal}(\beta_{\text{seas}}(t-1), \sigma) \qquad t = 2 \ldots 52 \tag{S95}$$

$$\beta_{\text{seas}}(1) \sim \text{Normal}(\beta_{\text{seas}}(52), \sigma) \tag{S96}$$

$$\sigma \sim \text{Half-Normal}(0, 1) \tag{S97}$$

We fixed $\alpha(t) = 1$ for all $t < 2020$ and estimate $\alpha$ assuming a normal prior:

$$\alpha \sim \text{Normal}(1, 0.25). \tag{S98}$$

We assumed weakly informative priors on $\omega$ and $\nu$:

$$\omega \sim \text{Half-Normal}(0, 200) \tag{S99}$$

$$\nu \sim \text{Normal}(104, 26) \tag{S100}$$

We assumed that the true birth/death rates, population sizes, and recovery rates are known. We note, however, that assuming $\gamma = 1/\text{week}$ actually corresponds to a mean simulated infectious period of 1.6 weeks due to a time discretization, which is much longer than the true value; this approximation allows us to test whether we can still robustly estimate the intrinsic resilience given parameter mis-specification. Initial conditions were estimated with following priors:

$$S(0) = Ns(0) \tag{S101}$$

$$I(0) = Ni(0) \tag{S102}$$

$$s(0) \sim \text{Uniform}(0, 1) \tag{S103}$$

$$i(0) \sim \text{Half-Normal}(0, 0.001) \tag{S104}$$

Finally, the observation model was specified as follows:

$$\text{Cases}(t) \sim \text{Negative-Binomial}(\rho N_{SI}(t), \phi) \tag{S105}$$

$$\rho \sim \text{Half-Normal}(0, 0.02) \tag{S106}$$

$$\phi \sim \text{Half-Normal}(0, 10) \tag{S107}$$

where $\rho$ represents the reporting probability and $\phi$ represents the negative binomial overdispersion parameter.

The model was fitted to four separate time series: (1) incidence time series from the SIRS model, (2) incidence time series for strain 1 from the two-strain model, (3) incidence time series for strain 2 from the two-strain model, and (4) combined incidence time series for strains 1 and 2 from the two-strain model. The model was fitted using rstan [6, 7] with 4 chains, each consisting of 2000 iterations. The resulting posterior distribution was used to calculate the intrinsic resilience of the seasonally unforced SIRS model with the same parameters; eigenvalues of the discrete-time SIR model were computed by numerically finding the equilibrium and calculating the Jacobian matrix.

### Validations for window-selection criteria

We used stochastic SIRS simulations to identify optimal parameters for the window-selection criteria that we used for the linear regression for estimating empirical resilience. For each simulation, we began by generating a random perturbation $\alpha(t)$ from a random set of parameters. First, we drew the duration of perturbation $\tau_{\text{npi}}$ from a uniform distribution between 1 and 2 years. Then, we drew independent normal variables $z_i$ of length $\lfloor 364\tau_{\text{npi}} \rfloor$ with a standard deviation of 0.02 and took a reverse cumulative sum to obtain a realistic shape for the perturbation:

$$x_n = 1 + \sum_{i=n}^{\lfloor 364\tau_{\text{npi}} \rfloor} z_i, \quad n = 1, \ldots, \lfloor 364\tau_{\text{npi}} \rfloor. \tag{S108}$$

In contrast to simple perturbations that assume a constant reduction in transmission, this approach allows us to model transmission reduction that varies over time smoothly. We repeated this random generation process until less than 10% of $x_n$ exceeds 1—this was done to ensure the perturbation term $\alpha(t)$ stays below 1 (and therefore reduce transmission) for the most part. Then, we set any values that are above 1 or below 0 to 1 and 0, respectively. Then, we randomly drew the minimum transmission during perturbation $\alpha_{\text{min}}$ from a uniform distribution between 0.5 and 0.7 and scale $x_n$ to have a minimum of $\alpha_{\text{min}}$:

$$x_{\text{scale},n} = \alpha_{\text{min}} + (1 - \alpha_{\text{min}}) \times \frac{x_n - \min x_n}{1 - \min x_n}. \tag{S109}$$

This allowed us to simulate a realistically shaped perturbation:

$$\alpha(t) = \begin{cases} 1 & t < 2020 \\ x_{\text{scale},364(t-2020)} & 2020 \le t < 2020 + \tau_{\text{npi}} \\ 1 & \tau_{\text{npi}} \le t \end{cases}. \tag{S110}$$

Given this perturbation function, we draw $\mathcal{R}_0$ from a uniform distribution between 1.5 and 4 and the mean duration of immunity $1/\delta$ from a uniform distribution between 1 and 4. Then, we simulate the stochastic SIRS model from $S(0) = 10^8/\mathcal{R}_0$ and $I(0) = 100$ from 1990 to 2025 and truncate the time series to 2014–2025; if the epidemic becomes extinct before the end of simulation, we discard that simulation and start over from the perturbation generation step.

For each epidemic simulation, we computed the empirical resilience by varying the threshold $R$ for the nearest neighbor approach from 4 to 14 with increments of 2, the number of divisions $K$ for the window selection between 8 and 25, and the truncation threshold $a$ for the window selection between 1 to 3; this was done for all possible combinations of $R$, $K$, and $a$. We also compared this with the naive approach that uses the entire distance-from-attractor time series, starting from the maximum distance to the end of the time series. We repeated this procedure 500 times and quantified the correlation between empirical and intrinsic resilience estimates across two approaches.

### Supplementary Results

#### Window-selection criteria and choices about embedding dimensions

We tested our automated window selection criterion against randomized, stochastic simulations across a range of realistic pandemic perturbation shapes. We also explored optimal choices for embedding dimensions and fitting window parameters by quantifying correlation coefficients between the estimated resilience and the intrinsic resilience of a seasonally unforced system (Materials and Methods). Estimation performances varied widely (correlation coefficients ranging from 0.21 to 0.61), but the automated approach consistently outperformed a naive approach, which relies on all data from peak distance to the present (Supplementary Figure S9).

We then applied the best-performing window selection criterion and embedding dimension to pathogen surveillance data (Materials and Methods). For each time series, we applied Takens' theorem independently to reconstruct the empirical attractor and obtained the corresponding time series of distances from attractors (Supplementary Figure S10). Then, we fitted a linear regression and estimated the empirical resilience for each pathogen in each country (Supplementary Figure S10). The window selection criterion gave poor regression windows in three cases (norovirus in Hong Kong and Korea and rhinovirus/enterovirus in the US), leading to unrealistically low resilience estimates, and so we used ad-hoc regression windows instead (Supplementary Figure S11; Materials and Methods).

#### Out-of-sample predictions

This led to greater uncertainty in resilience estimates and corresponding predictions of the expected return time (Supplementary Figure S16A,B). The expected and observed return times exhibit weak positive correlation ($\rho = 0.36$; 95% CI: -0.34–0.81), and 7 out of 10 confidence intervals contain the true value (Supplementary Figure S16C). We found moderate correlations ($\rho = 0.51$; 95% CI: 0.01–0.80) in predictions for the expected return time between partial regression fits, which rely on observations up to the 26th week of 2023, and full regression fits, which rely on the entire data set (Supplementary Figure S16D); this implies consistency in our predictions. However, resilience estimates from partial regression fits and full regression fits were weakly correlated ($\rho = 0.30$; 95% CI: -0.23–0.70), demonstrating challenges in estimating pathogen resilience from limited data (Supplementary Figure S16E).

### Supplementary Figures

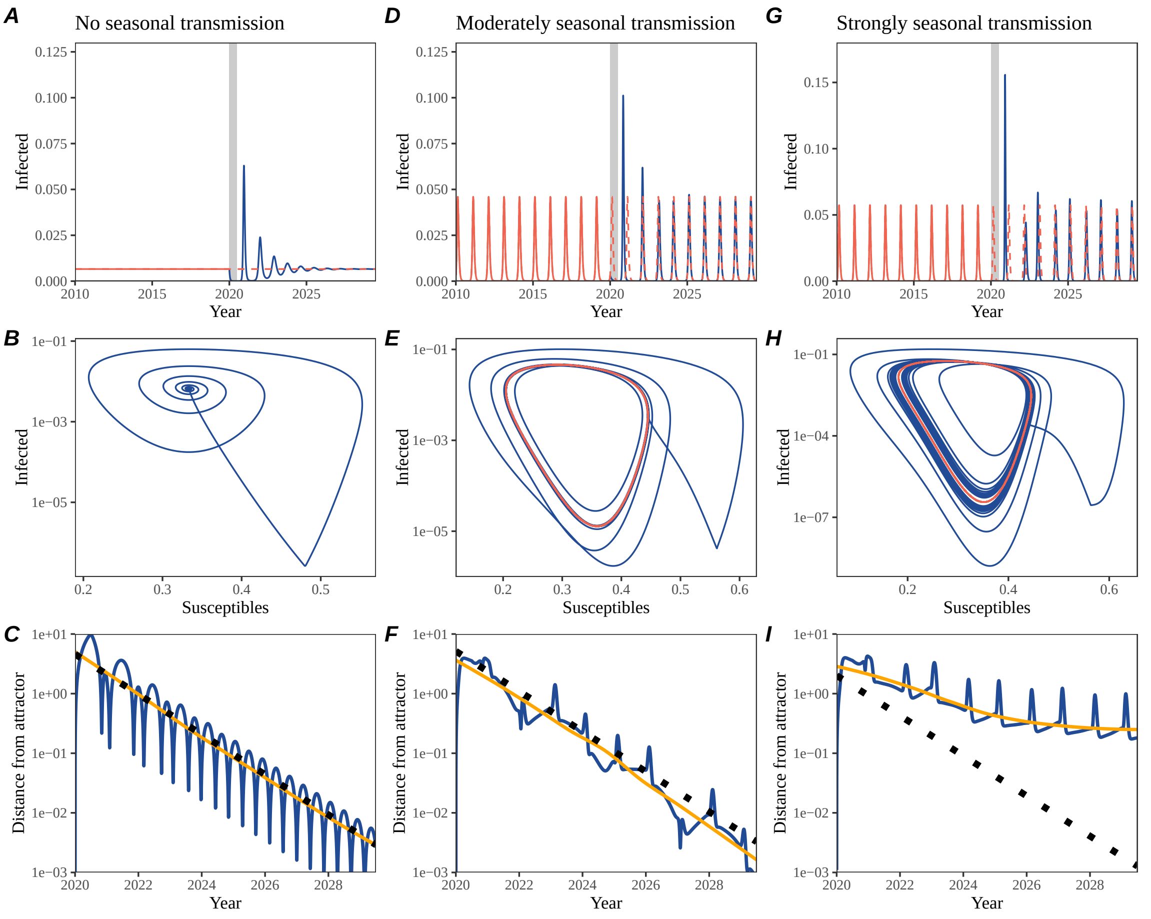


Figure S1: **Impact of seasonal transmission on pathogen resilience.** (A, D, G) Simulated epidemic trajectories using the SIRS model without seasonal forcing (A), with seasonal forcing of amplitude of 0.2 (D), and with seasonal forcing of amplitude of 0.4 (G). Red and blue solid lines represent epidemic dynamics before and after interventions are introduced, respectively. Red dashed lines represent counterfactual epidemic dynamics in the absence of interventions. Gray regions indicate the duration of interventions. (B, E, H) Phase plane representation of the time series in panels A, D, and G alongside the corresponding susceptible host dynamics. Red and blue solid lines represent epidemic trajectories on an SI phase plane before and after interventions are introduced, respectively. (C, F, I) Changes in logged distance from the attractor over time. Blue lines represent the logged distance from the attractor. Orange lines represent the locally estimated scatterplot smoothing (LOESS) fits to the logged distance from the attractor. Dotted lines show the intrinsic resilience of the seasonally unforced system.

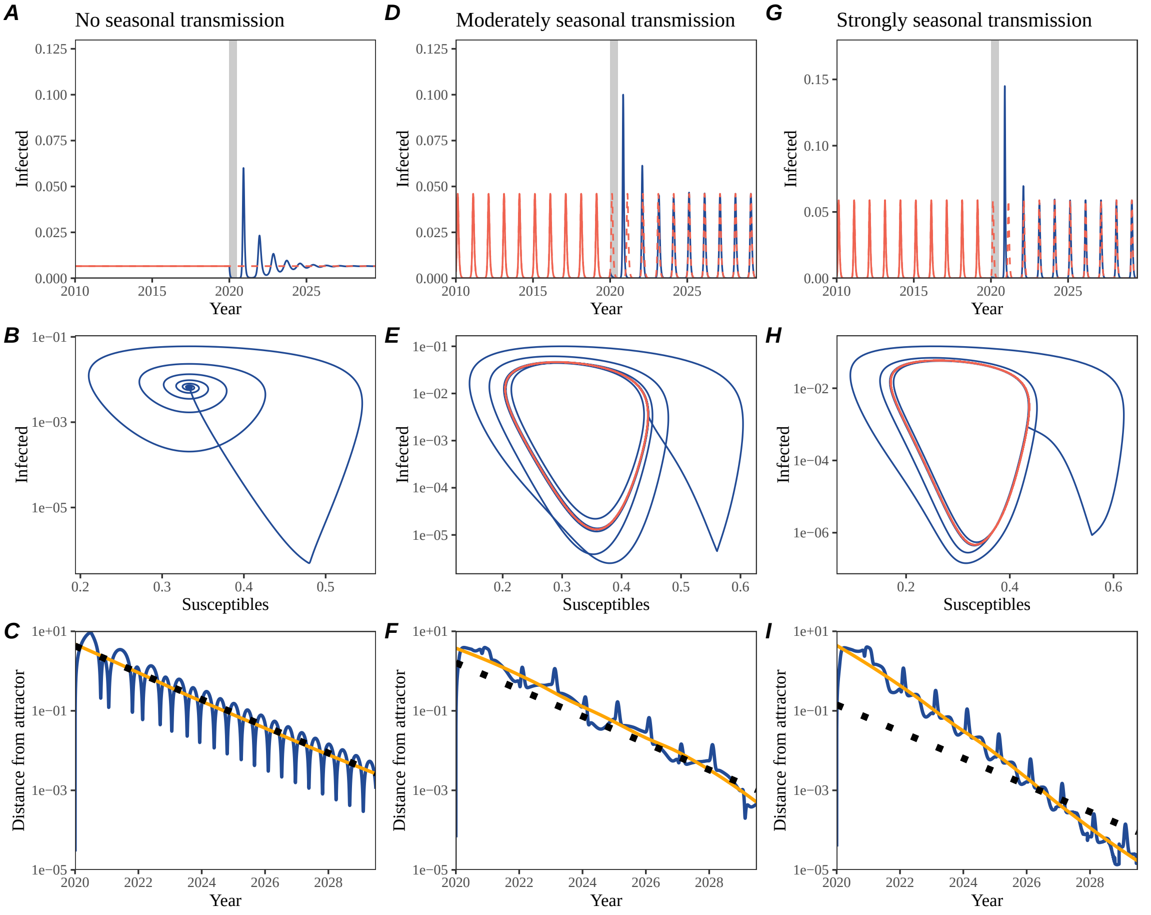


Figure S2: **Impact of seasonal transmission and importation on pathogen resilience.** (A, D, G) Simulated epidemic trajectories using the SIRS model without seasonal forcing (A), with seasonal forcing of amplitude of 0.2 (D), and with seasonal forcing of amplitude of 0.4 (G). Red and blue solid lines represent epidemic dynamics before and after interventions are introduced, respectively. Red dashed lines represent counterfactual epidemic dynamics in the absence of interventions. Gray regions indicate the duration of interventions. (B, E, H) Phase plane representation of the time series in panels A, D, and G alongside the corresponding susceptible host dynamics. Red and blue solid lines represent epidemic trajectories on an SI phase plane before and after interventions are introduced, respectively. (C, F, I) Changes in logged distance from the attractor over time. Blue lines represent the logged distance from the attractor. Orange lines represent the locally estimated scatterplot smoothing (LOESS) fits to the logged distance from the attractor. Dotted lines show the intrinsic resilience of the seasonally unforced system. A low amount of importation ($10^{-7}$ per day) is assumed for all scenarios.

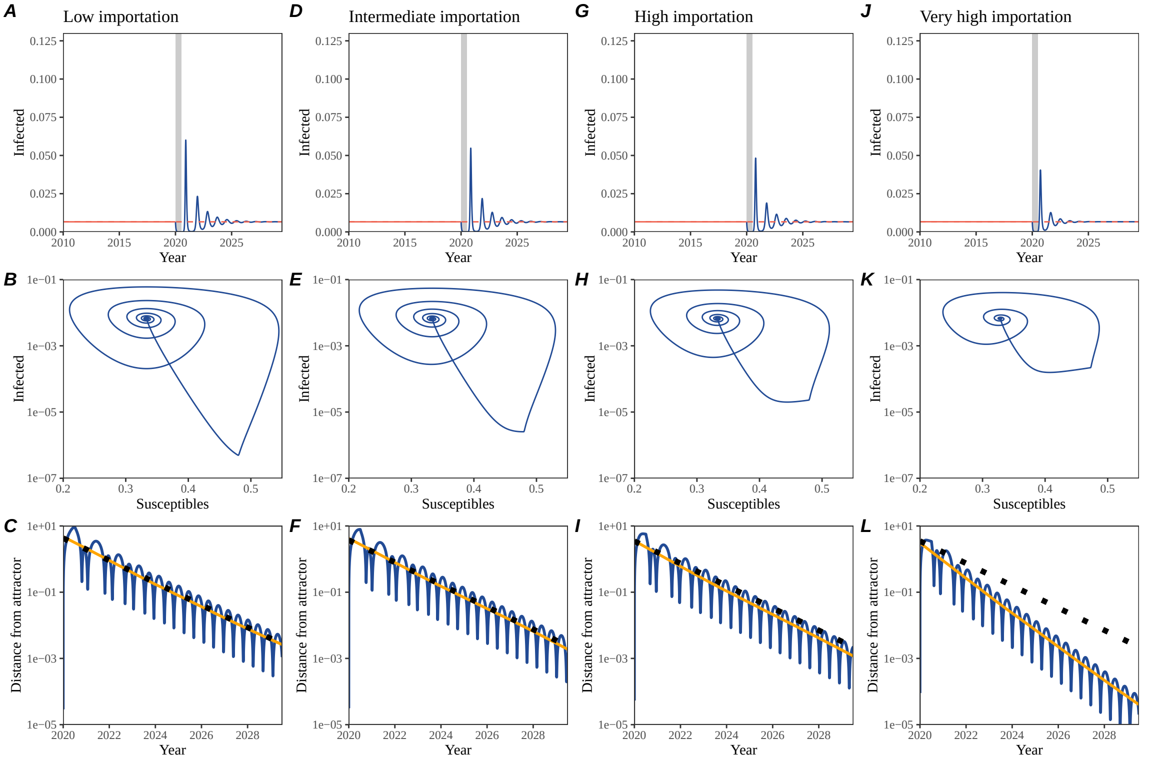


Figure S3: **Impact of importation on pathogen resilience.** (A, D, G, J) Simulated epidemic trajectories using the SIRS model without seasonal forcing assuming a low ($10^{-7}$ per day), moderate ($10^{-6}$ per day), high ($10^{-5}$ per day), and very high ($10^{-4}$ per day) amounts of importation. Red and blue solid lines represent epidemic dynamics before and after interventions are introduced, respectively. Red dashed lines represent counterfactual epidemic dynamics in the absence of interventions. Gray regions indicate the duration of interventions. (B, E, H, K) Phase plane representation of the time series in panels A, D, G, and J alongside the corresponding susceptible host dynamics. Red and blue solid lines represent epidemic trajectories on an SI phase plane before and after interventions are introduced, respectively. (C, F, I, L) Changes in logged distance from the attractor over time. Blue lines represent the logged distance from the attractor. Orange lines represent the locally estimated scatterplot smoothing (LOESS) fits to the logged distance from the attractor. Dotted lines show the intrinsic resilience of the seasonally unforced system.

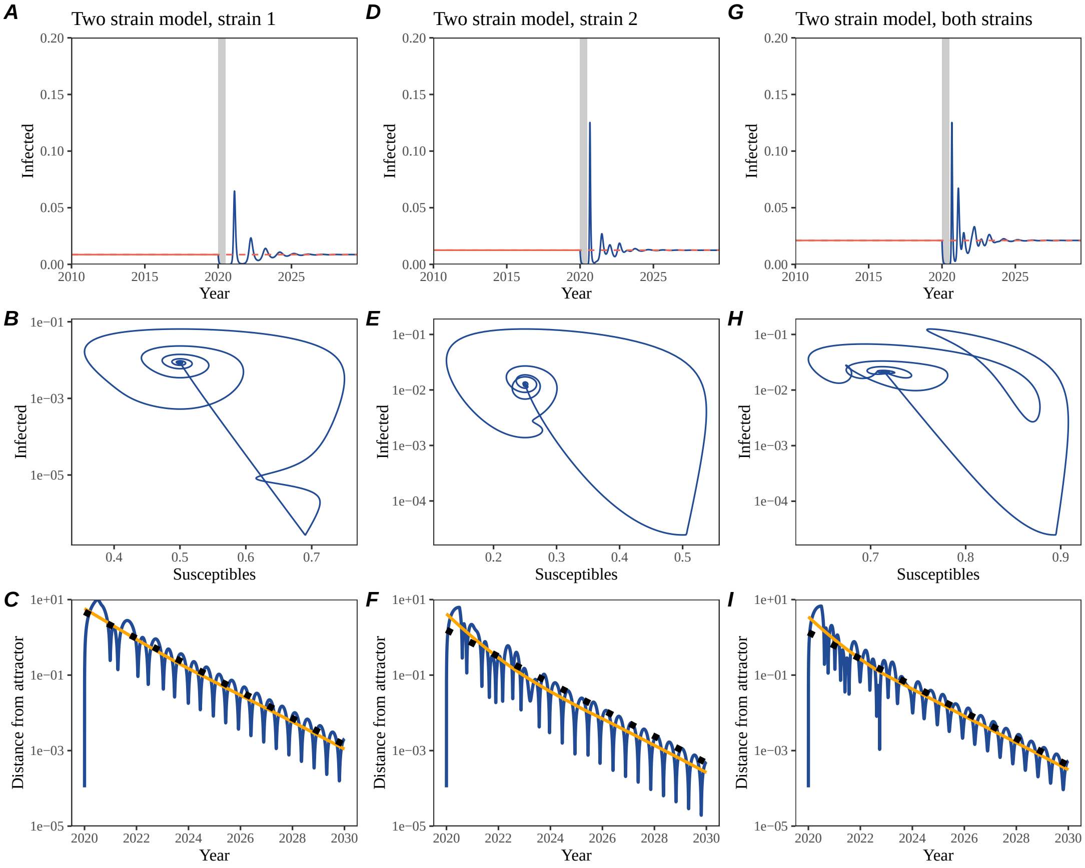


Figure S4: **A simple method to measure pathogen resilience following pandemic perturbations for a two-strain competition system without seasonal forcing.** (A, D, G) Simulated epidemic trajectories using a two-strain system without seasonal forcing. In this model, two strains compete through cross immunity. Red and blue solid lines represent epidemic dynamics before and after interventions are introduced, respectively. Red dashed lines represent counterfactual epidemic dynamics in the absence of interventions. Gray regions indicate the duration of interventions. (B, E, H) Phase plane representation of the time series in panels A, D, and G alongside the corresponding susceptible host dynamics. For the dynamics of strains 1 and 2, the susceptibles (x-axis) measure the mean susceptibility against each strain. For the joint dynamics, the susceptibles simply measure the total number of individuals who can be infected by either strains (Supplementary Text). Blue solid lines represent epidemic trajectories on an SI phase plane before and after interventions are introduced, respectively. (C, F, I) Changes in logged distance from the attractor over time. Blue lines represent the logged distance from the attractor. Orange lines represent the locally estimated scatterplot smoothing (LOESS) fits to the logged distance from the attractor. Dotted lines show the intrinsic resilience of the seasonally unforced system.

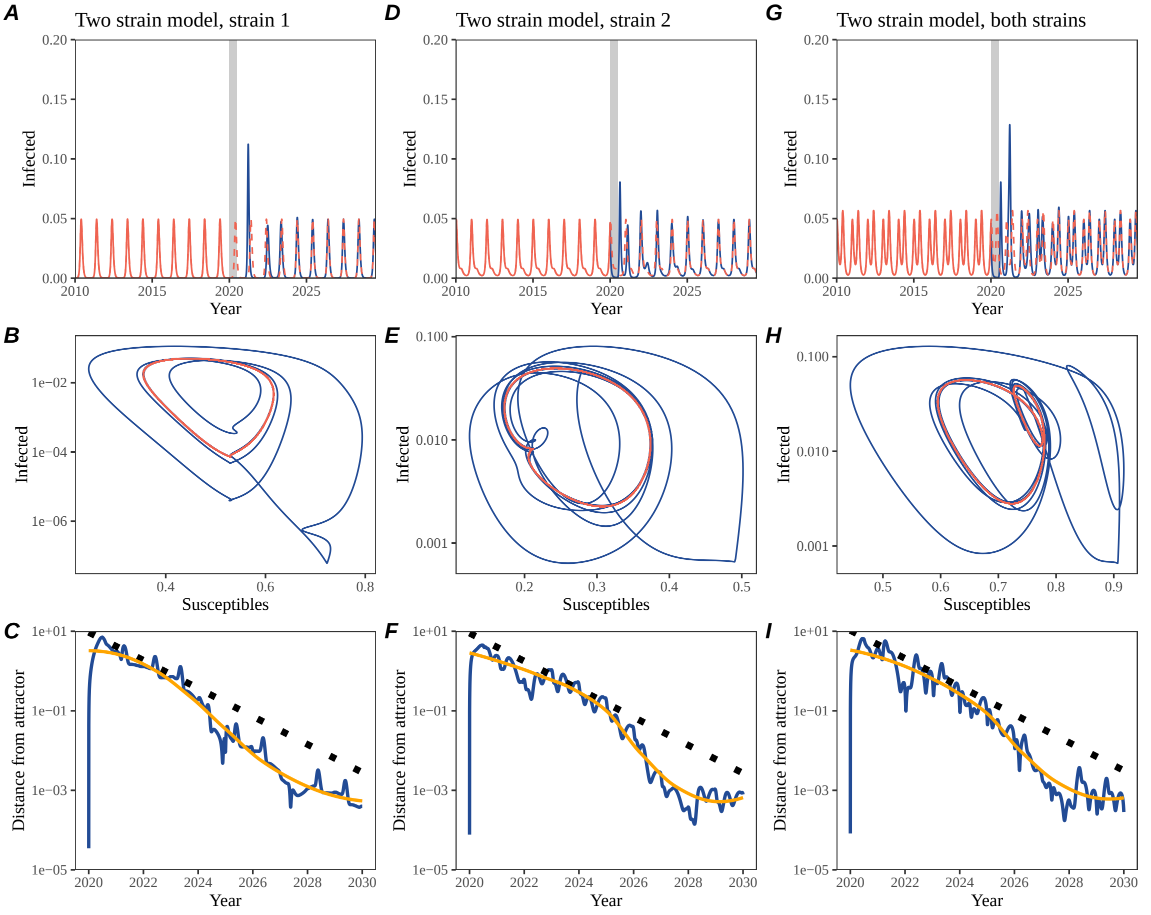


Figure S5: **A simple method to measure pathogen resilience following pandemic perturbations for a two-strain competition system with seasonal forcing.** (A, D, G) Simulated epidemic trajectories using a two-strain system with seasonal forcing (amplitude of 0.2). In this model, two strains compete through cross immunity. Red and blue solid lines represent epidemic dynamics before and after interventions are introduced, respectively. Red dashed lines represent counterfactual epidemic dynamics in the absence of interventions. Gray regions indicate the duration of interventions. (B, E, H) Phase plane representation of the time series in panels A, D, and G alongside the corresponding susceptible host dynamics. Red and blue solid lines represent epidemic trajectories on an SI phase plane before and after interventions are introduced, respectively. (C, F, I) Changes in logged distance from the attractor over time. Blue lines represent the logged distance from the attractor. Orange lines represent the locally estimated scatterplot smoothing (LOESS) fits to the logged distance from the attractor. Dotted lines show the intrinsic resilience of the seasonally unforced system.

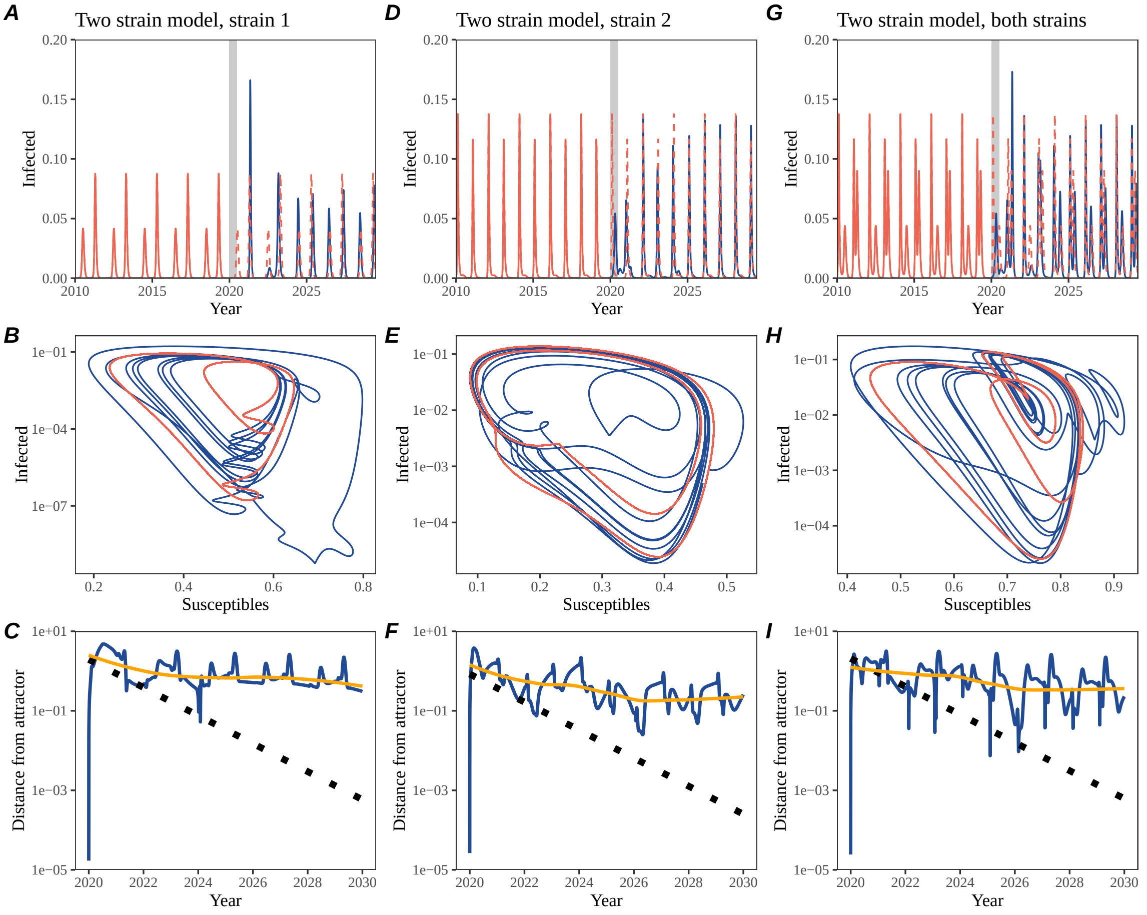


Figure S6: **A simple method to measure pathogen resilience following pandemic perturbations for a multi-strain competition system with strong seasonal forcing.** (A, D, G) Simulated epidemic trajectories using a two-strain system with seasonal forcing (amplitude of 0.2). In this model, two strains compete through cross immunity. Red and blue solid lines represent epidemic dynamics before and after interventions are introduced, respectively. Red dashed lines represent counterfactual epidemic dynamics in the absence of interventions. Gray regions indicate the duration of interventions. (B, E, H) Phase plane representation of the time series in panels A, D, and G alongside the corresponding susceptible host dynamics. Red and blue solid lines represent epidemic trajectories on an SI phase plane before and after interventions are introduced, respectively. (C, F, I) Changes in logged distance from the attractor over time. Blue lines represent the logged distance from the attractor. Orange lines represent the locally estimated scatterplot smoothing (LOESS) fits to the logged distance from the attractor. Dotted lines show the intrinsic resilience of the seasonally unforced system.

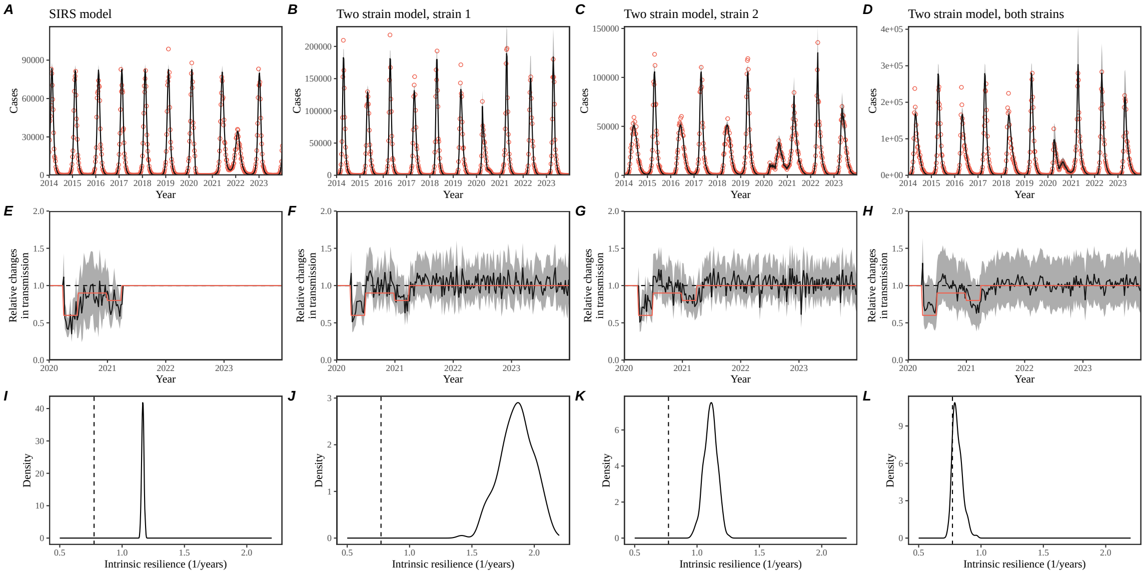


Figure S7: **Mechanistic model fits to simulated data and inferred intrinsic resilience.** We simulated discrete time, stochastic epidemic trajectories using a seasonally forced SIRS model (A,E,I) and a seasonally forced two-strain model (B–D, F–H, and J–L) and tested whether we can estimate the intrinsic resilience of the seasonally unforced versions of the model by fitting a mechanistic model (Supplementary Text). We fitted the same discrete time, one strain, deterministic SIRS model in each scenario. (A–D) Simulated case time series from corresponding models shown in the title (red) and SIRS model fits (black). (E–H) Assumed changes in transmission due to pandemic perturbations (red) and estimated time-varying relative transmission rates for the perturbation impact from the SIRS model fits (black). Solid lines and shaded regions represent fitted posterior median and 95% credible intervals. (I–L) Comparisons between the true and estimated intrinsic resilience of the seasonally unforced system. Vertical lines represent the true intrinsic resilience of the seasonally unforced system. Density plots represent the posterior distribution of the inferred intrinsic resilience of the seasonally unforced SIRS model using fitted parameters.

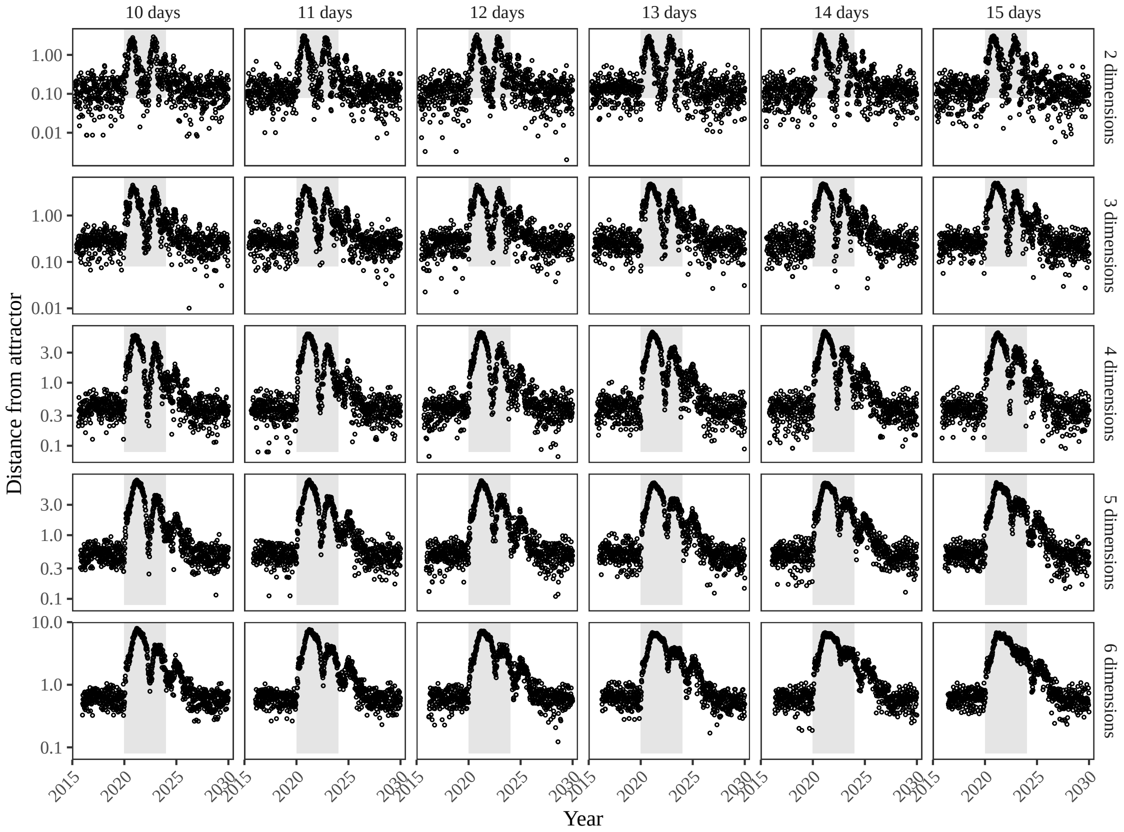


Figure S8: **Sensitivity of the distance from the attractor to choice of embedding lags and dimensions.** Sensitivity analysis for the distance-from-attractor time series shown in Figure 3E in the main text by varying the embedding lag between 10–15 days and embedding dimensions between 2–6 dimensions. The gray region represents the assumed period of pandemic perturbation.

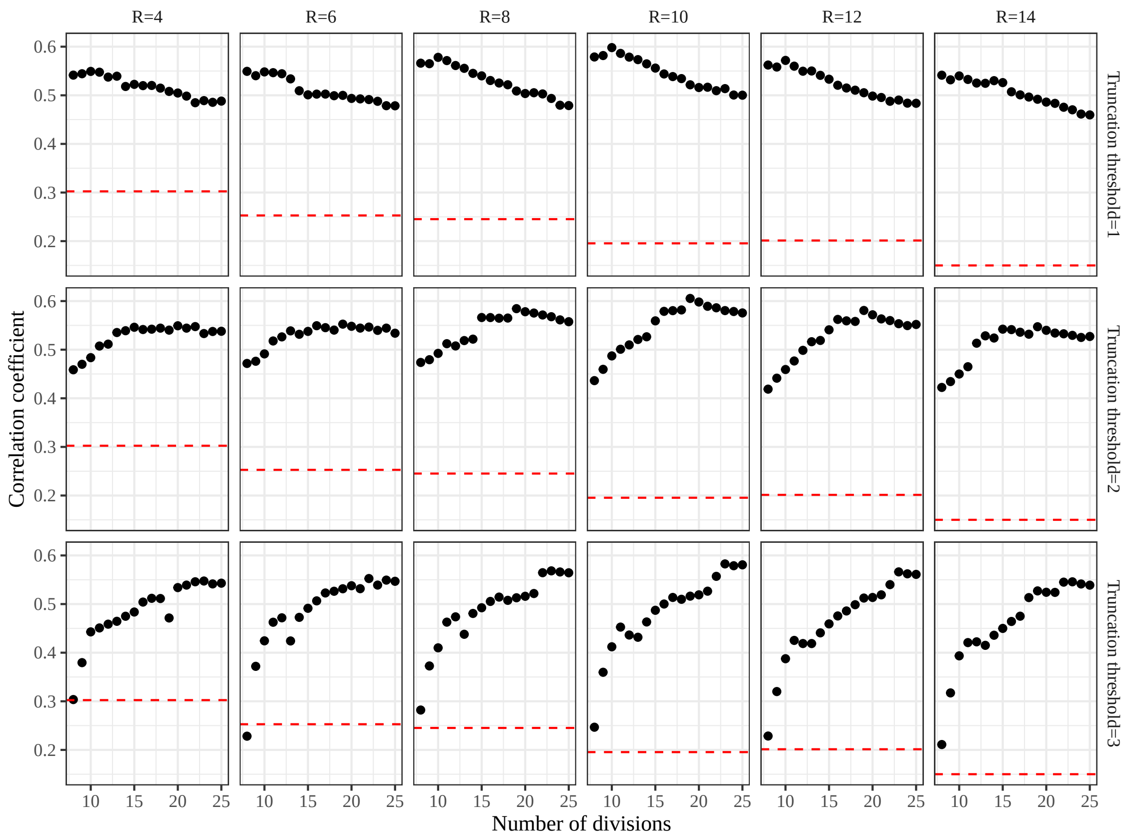


Figure S9: **Impact of fitting window selection on the estimation of empirical resilience.** We simulated 500 epidemics of a stochastic SIRS model with randomly drawn parameters and randomly generated pandemic perturbations (Supplementary Text). For each simulation, we varied the false nearest neighbor threshold $R$ for reconstructing the empirical attractor as well as the parameters for regression window selection (i.e., the truncation threshold $a$ and the number of divisions $K$). Each point represents the resulting correlation coefficient between empirical and intrinsic resilience estimates for a given scenario. Dashed lines represent the correlation coefficient between empirical and intrinsic resilience estimates for the naive approach that fits a linear regression on a log scale, starting from the maximum distance until the end of the time series.

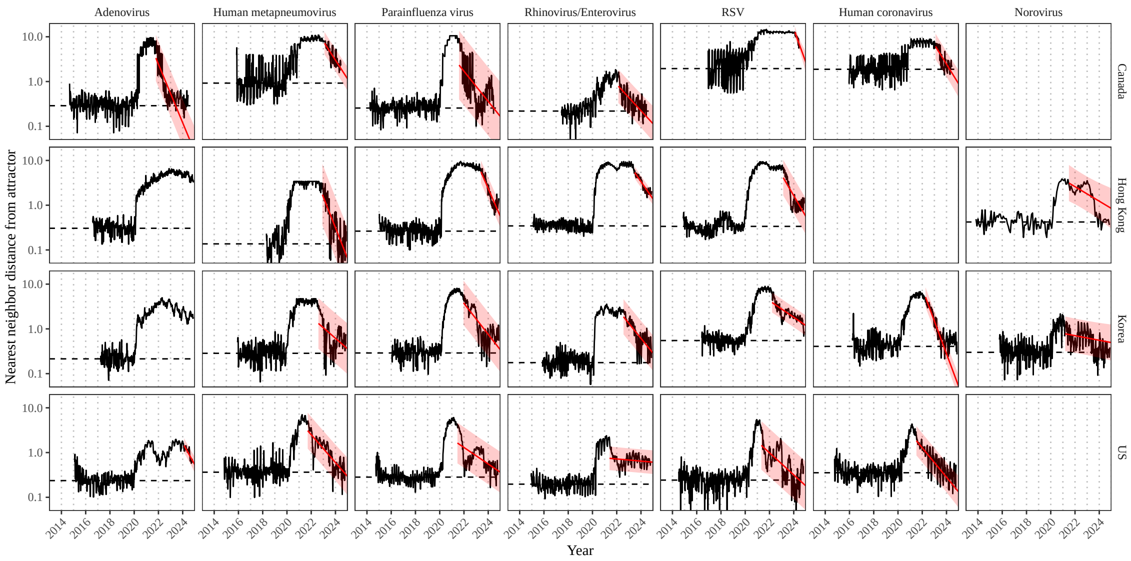


Figure S10: **Estimated time series of distance from the attractor for each pathogen and corresponding linear regression fits using automated window selection criteria across Canada, Hong Kong, Korea, and the US.** Black lines represent the estimated distance from the attractor. Red lines and shaded regions represent the linear regression fits and corresponding 95% confidence intervals. Dashed lines represent the average of the pre-pandemic nearest neighbor distances.

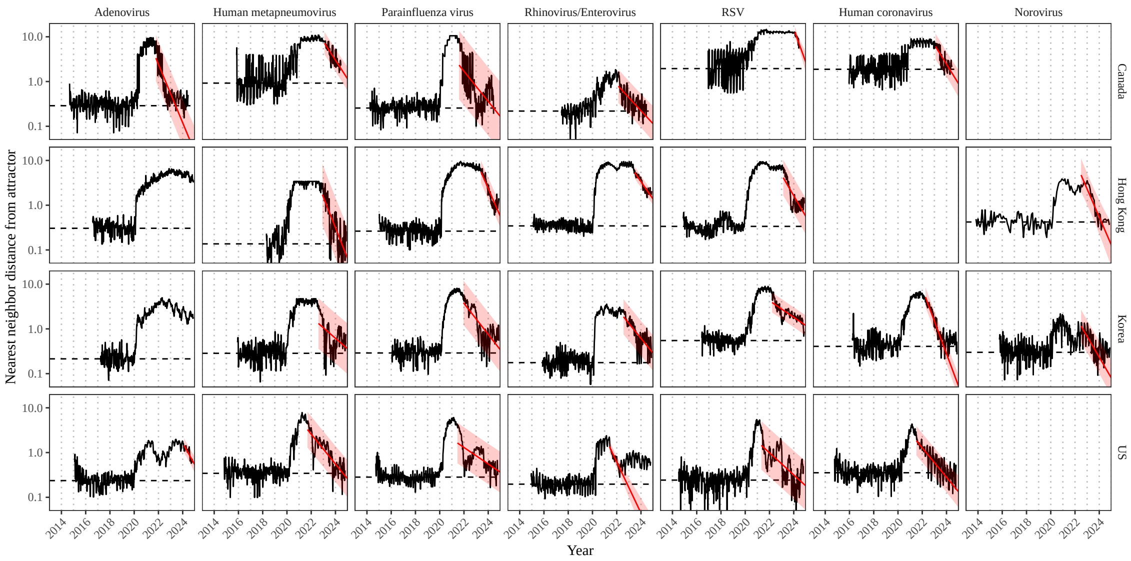


Figure S11: **Estimated time series of distance from the attractor for each pathogen and corresponding linear regression fits across Canada, Hong Kong, Korea, and the US, including ad-hoc regression window selection.** We used ad-hoc regression windows for norovirus in Hong Kong and Korea and rhinovirus/enterovirus in the US. Black lines represent the estimated distance from the attractor. Red lines and shaded regions represent the linear regression fits and corresponding 95% confidence intervals. Dashed lines represent the average of the pre-pandemic nearest neighbor distances.

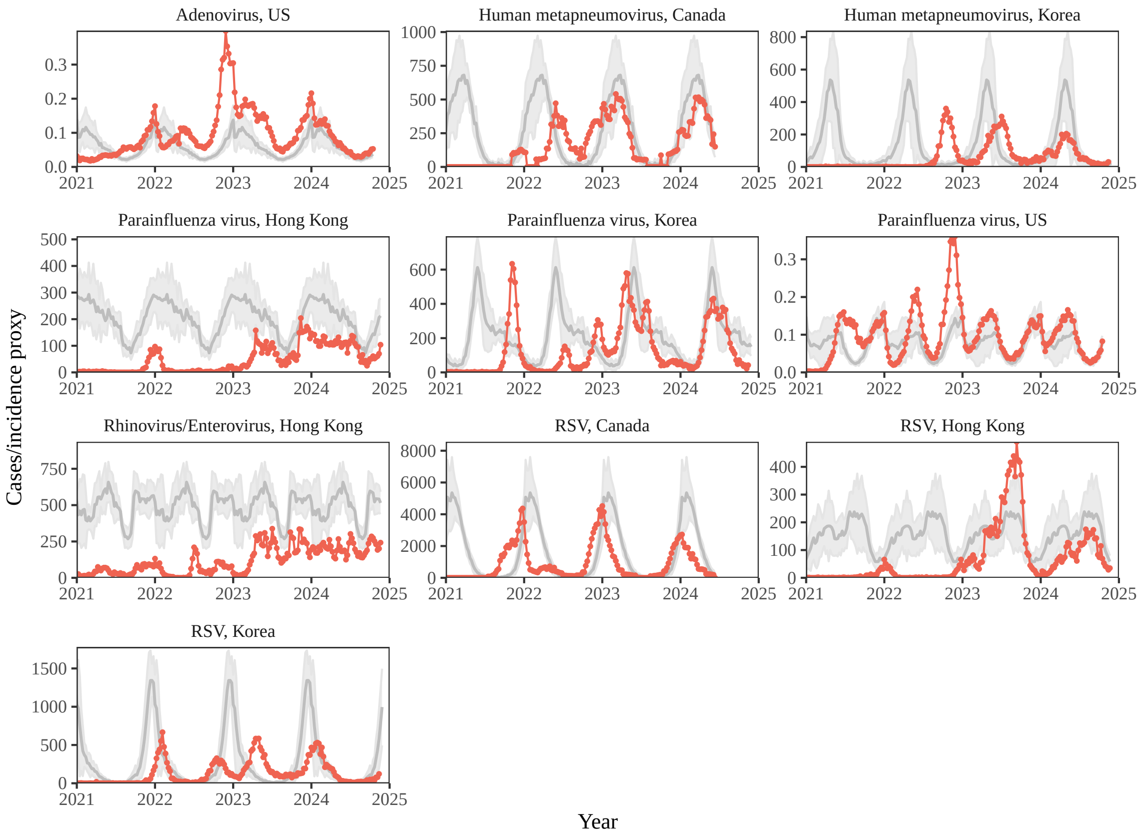


Figure S12: **Observed dynamics for pathogen that are predicted to return after the end of 2024.** Red points and lines represent data before 2020. Gray lines and shaded regions represent the mean seasonal patterns and corresponding 95% confidence intervals around the mean, previously shown in Figure 1.

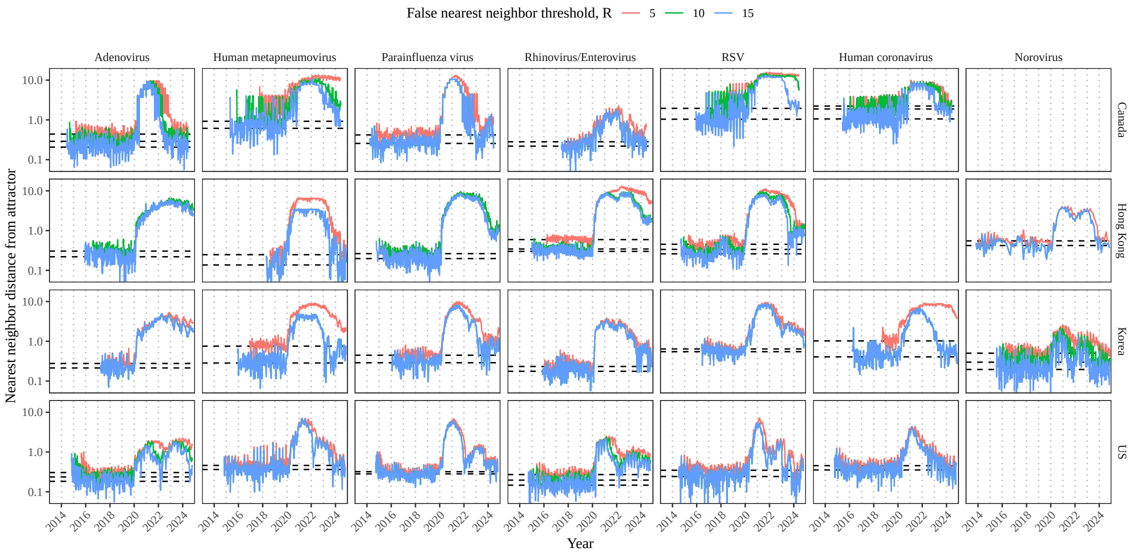


Figure S13: **Estimated time series of distance from the attractor for each pathogen across different choices about false nearest neighbor threshold values.** Using higher threshold values for the false nearest neighbor approach gives lower embedding dimensions. Colored lines represent the estimated distance from the attractor for different threshold values.

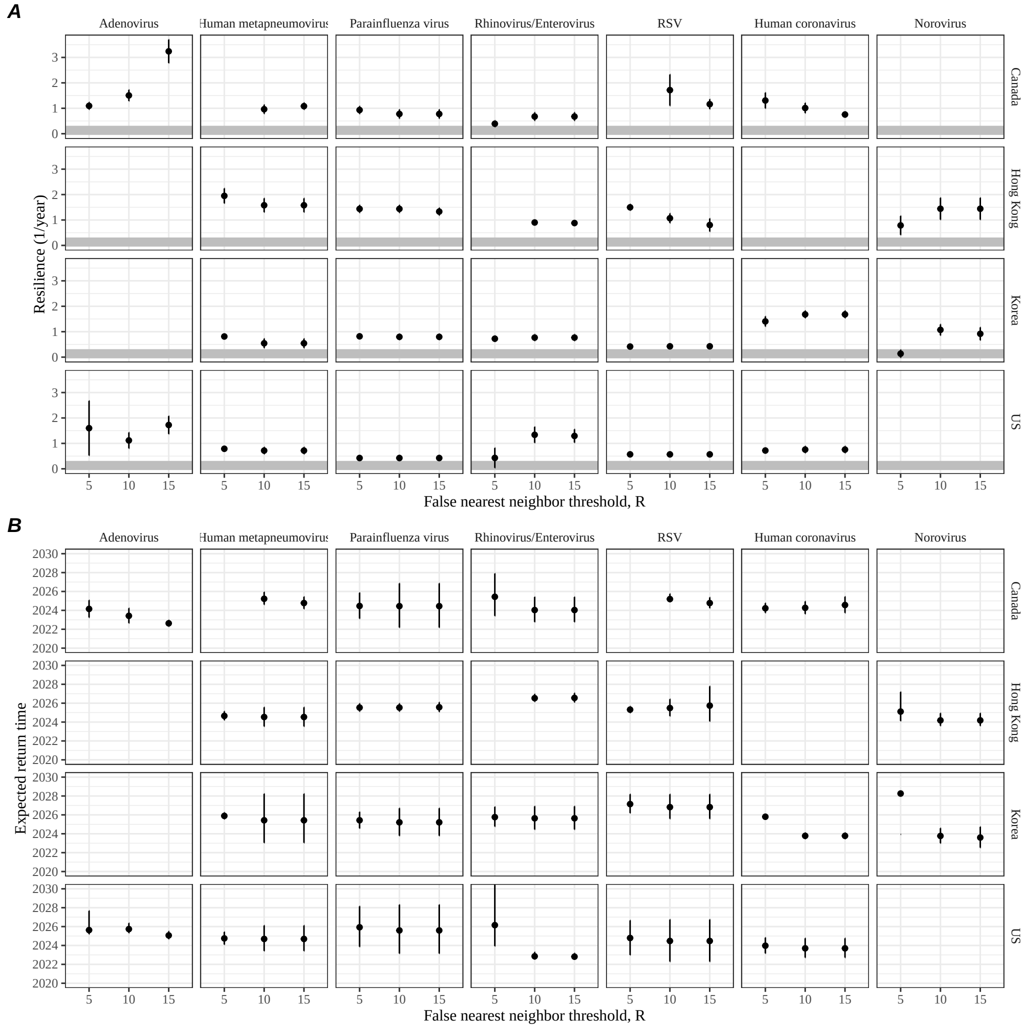


Figure S14: **Summary of resilience estimates and predictions for return time across different choices about false nearest neighbor threshold values.** The automated window selection method was used for all scenarios to estimate pathogen resilience, except for norovirus in Hong Kong and Korea and rhinovirus/enterovirus in the US. We used the same ad-hoc regression windows for norovirus in Hong Kong and Korea and rhinovirus/enterovirus in the US as we did in the main analysis. (A) Estimated pathogen resilience. The gray horizontal line represents the intrinsic resilience of pre-vaccination measles dynamics. (B) Predicted timing of when each pathogen will return to their pre-pandemic cycles. The dashed line in panel B indicates the end of 2024. Error bars represent 95% confidence intervals.

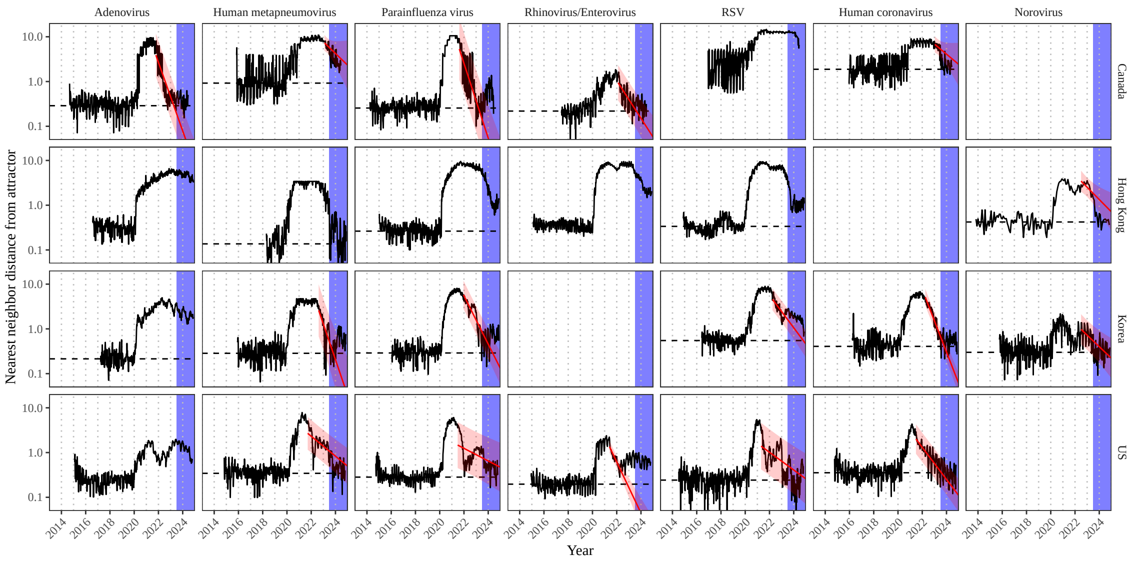


Figure S15: **Estimated time series of distance from the attractor for each pathogen and corresponding linear regression fits across Canada, Hong Kong, Korea, and the US based on data up to the 26th week of 2023.** We limited our data up to the 26th week of 2023 and applied the same regression approach as before. We used the same ad-hoc regression windows for norovirus in Hong Kong and Korea and rhinovirus/enterovirus in the US as the main analysis. Blue regions represent out of sample data. Black lines represent the estimated distance from the attractor. Red lines and shaded regions represent the linear regression fits and corresponding 95% confidence intervals. Dashed lines represent the average of the pre-pandemic nearest neighbor distances.

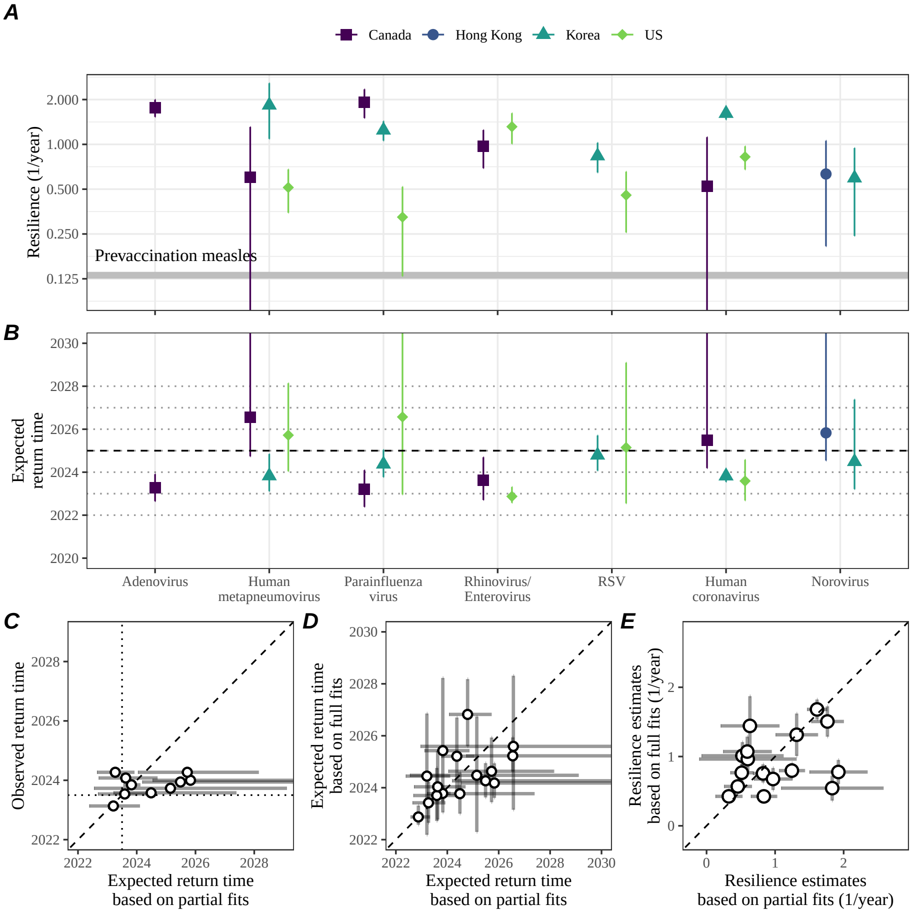


Figure S16: **Summary of resilience estimates and out-of-sample predictions for return time based on data up to the 26th week of 2023.** (A) Estimated pathogen resilience. The gray horizontal line represents the intrinsic resilience of pre-vaccination measles dynamics. (B) Predicted timing of each pathogen's return to their pre-pandemic cycles. The dashed line in panel B indicates the end of 2024. Error bars represent 95% confidence intervals. (C) Comparisons between the predicted and observed return time. Vertical and horizontal dotted lines represent the 26th week of 2023; therefore, all predictions for points above this horizontal lines represent out-of-sample predictions. (D) Comparisons between predicted return times based on partial fits (based on data up to the 26th week of 2023) and full fits (based on all data). (E) Comparisons between resilience estimates based on partial fits (based on data up to the 26th week of 2023) and full fits (based on all data). Dashed lines represent the one-to-one line. Vertical and horizontal dotted lines

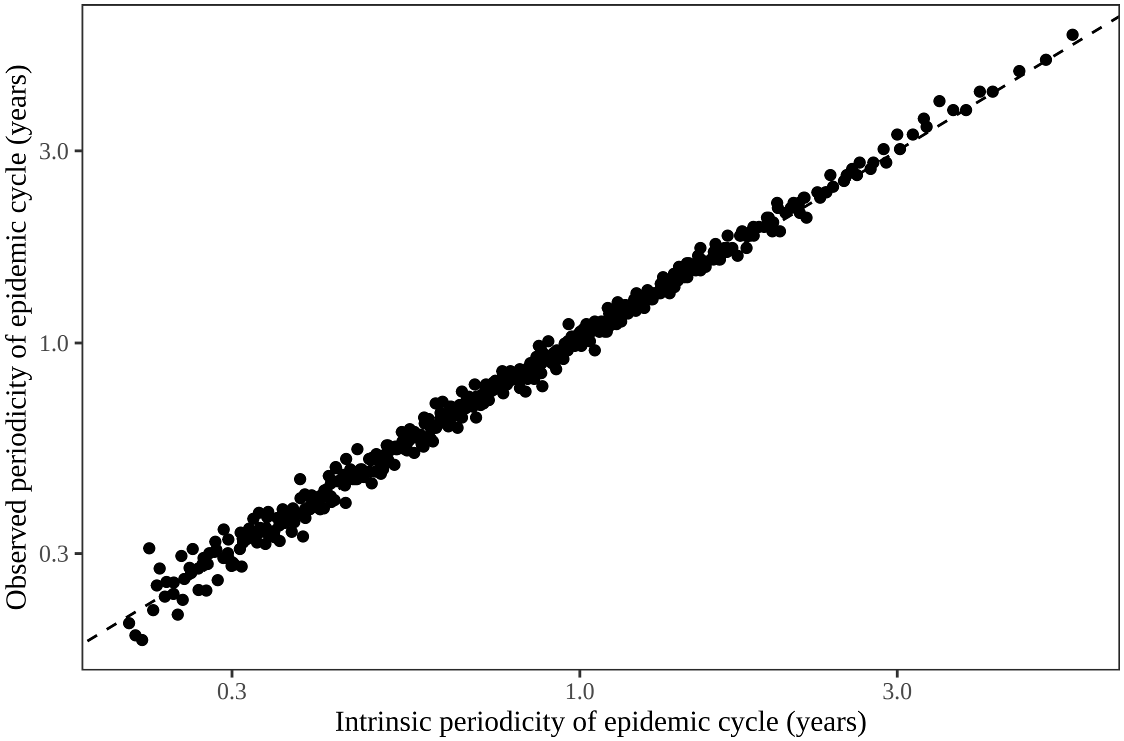


Figure S17: **Comparison between the observed and intrisic periodicity of the epidemic cycle of the seasonally unforced SIRS model.** The observed periodicity of the epidemic corresponds to the periodicity at which maximum spectral density occurs. The intrinsic periodicity of the epidemic corresponds to $2\pi/\text{Im}(\lambda)$, where $\text{Im}(\lambda)$ is the imaginary part of the eigenvalue.

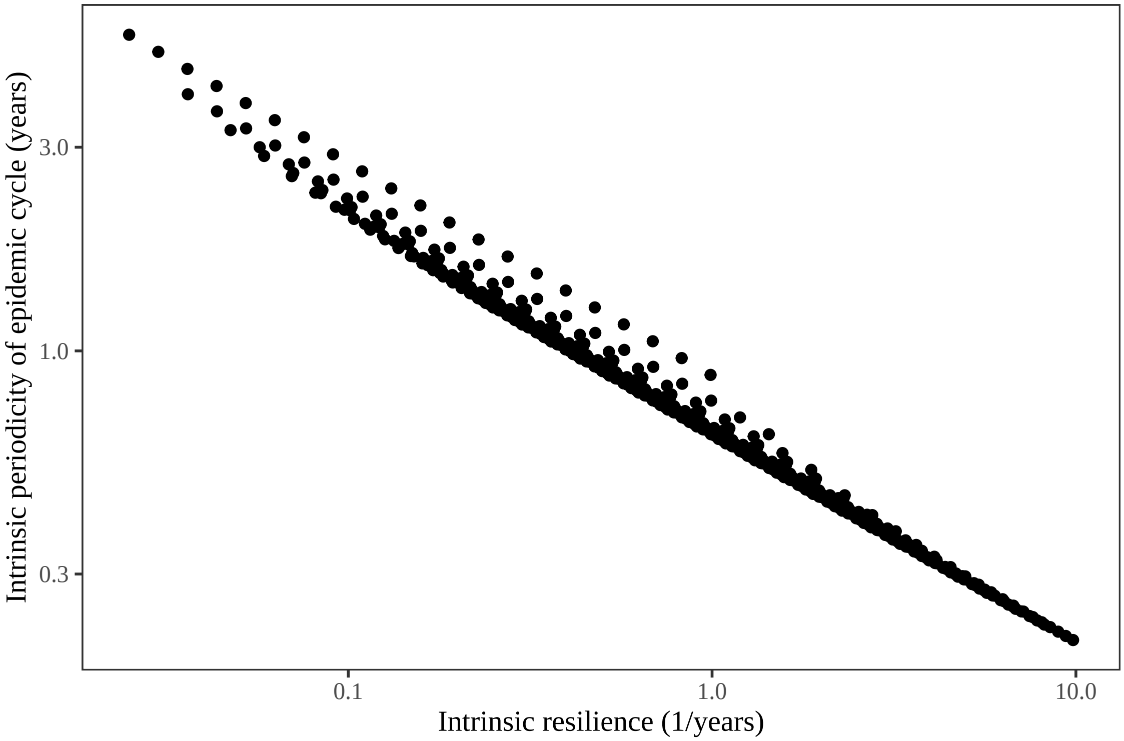


Figure S18: **Comparison between the intrisic resilience and periodicity of the epidemic cycle of the seasonally unforced SIRS model.** The intrinsic resilience of the epidemic corresponds to the negative of the real value of the eigenvalue $-\mathrm{Re}(\lambda)$. The intrinsic periodicity of the epidemic corresponds to $2\pi/\mathrm{Im}(\lambda)$, where $\mathrm{Im}(\lambda)$ is the imaginary part of the eigenvalue.

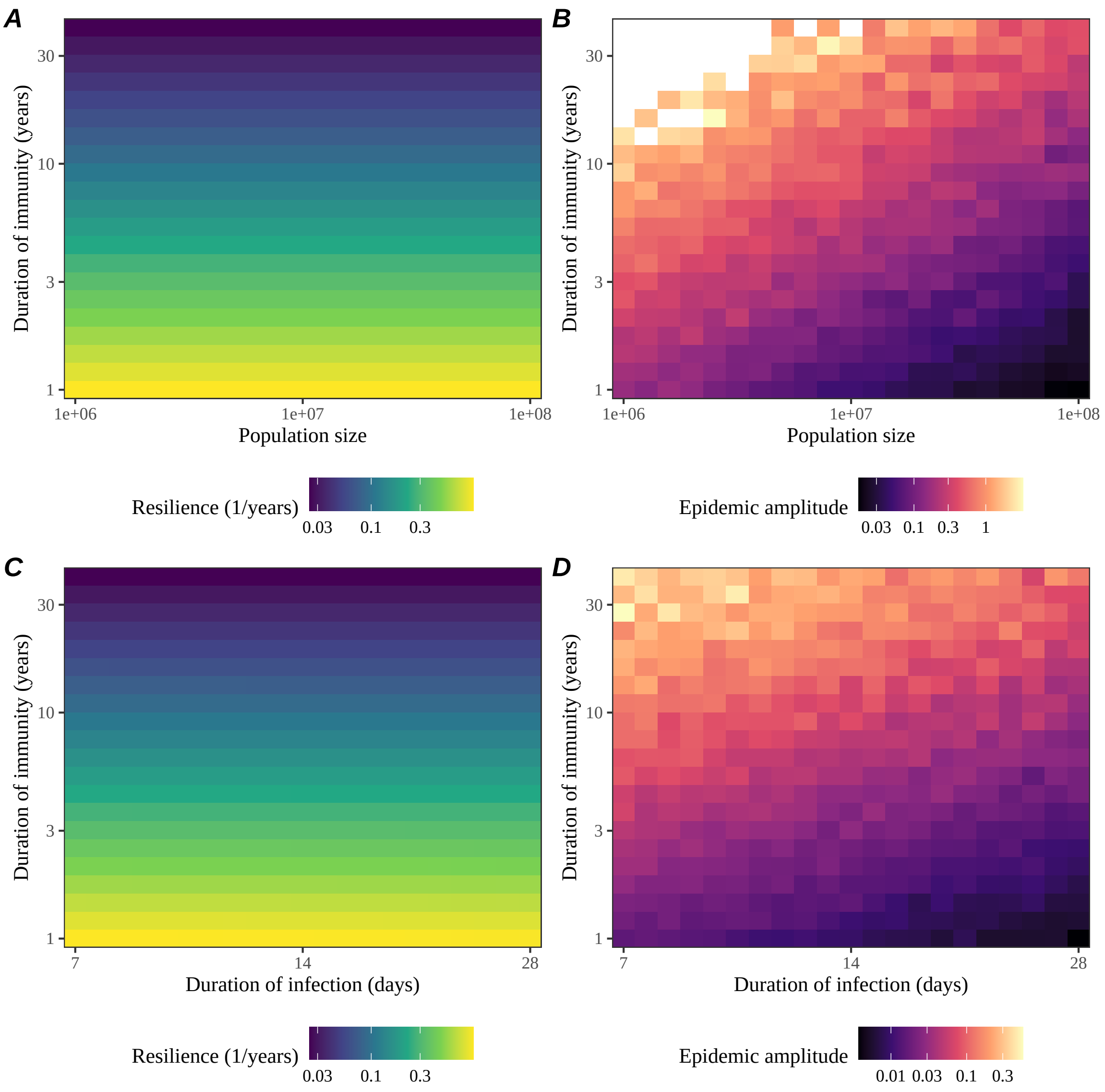


Figure S19: **Impact of population size and the average duration of infection of a host-pathogen system on its sensitivity to stochastic perturbations.** (A) Intrinsic resilience of a system as a function of population size and the average duration of immunity. (B) Epidemic amplitude as a function of population size and the average duration of immunity. (C) Intrinsic resilience of a system as a function of the average duration of infection and immunity. (D) Epidemic amplitude as a function of the average duration of infection and immunity. The epidemic amplitude corresponds to $(\max I - \min I)/(2\bar{I})$, where $\bar{I}$ represents the mean prevalence.

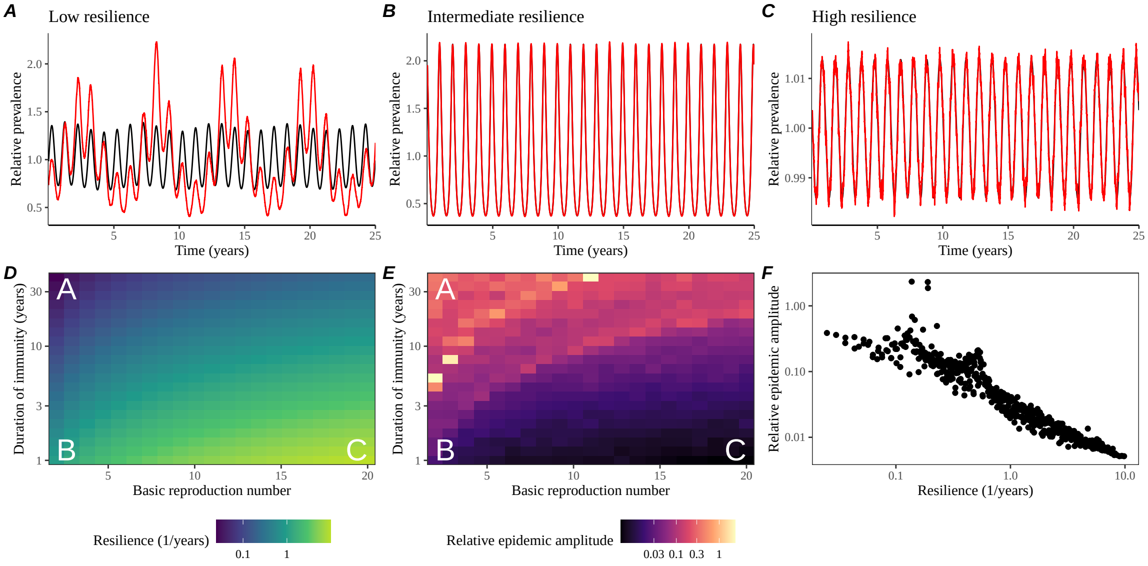


Figure S20: **Linking resilience of a host-pathogen system to its sensitivity to stochastic perturbations for the seasonally forced SIRS model.** (A–C) Epidemic trajectories of a stochastic SIRS model with demographic stochasticity across three different resilience values: low, intermediate, and high. The relative prevalence was calculated by dividing infection prevalence by its mean value. Black lines represent deterministic epidemic trajectories. Red lines represent stochastic epidemic trajectories. (D) The intrinsic resilience of the seasonally unforced system as a function of the basic reproduction number $\mathcal{R}_0$ and the duration of immunity. (E) The relative epidemic amplitude, a measure of sensitivity to stochastic perturbations, as a function of the basic reproduction number $\mathcal{R}_0$ and the duration of immunity. To calculate the relative epidemic amplitude, we first take the relative difference in infection prevalence between stochastic and deterministic trajectories: $\epsilon = (I_{\text{stoch}} - I_{\text{det}})/I_{\text{det}}$. Then, we calculate the difference between maximum and minimum of the relative difference and divide by half: $(\max \epsilon - \min \epsilon)/2$. Labels A–C in panels D and E correspond to scenarios shown in panels A–C. (F) The relationship between pathogen resilience and relative epidemic amplitude.

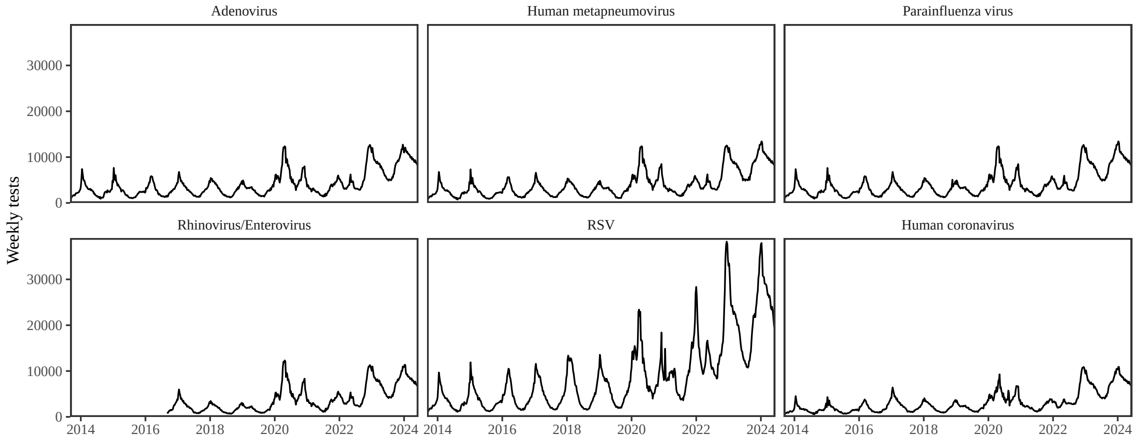


Figure S21: **Testing patterns for respiratory pathogens in Canada.**

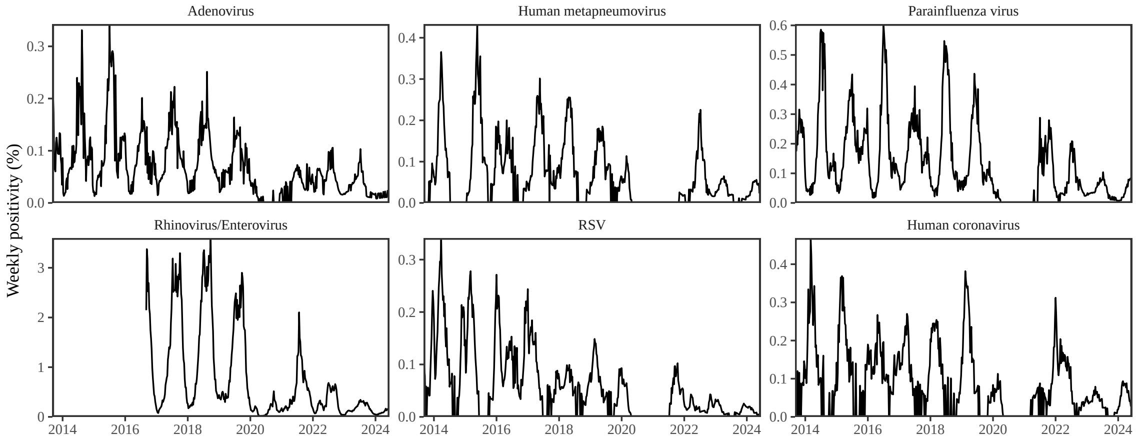


Figure S22: **Test positivity patterns for respiratory pathogens in Canada.**

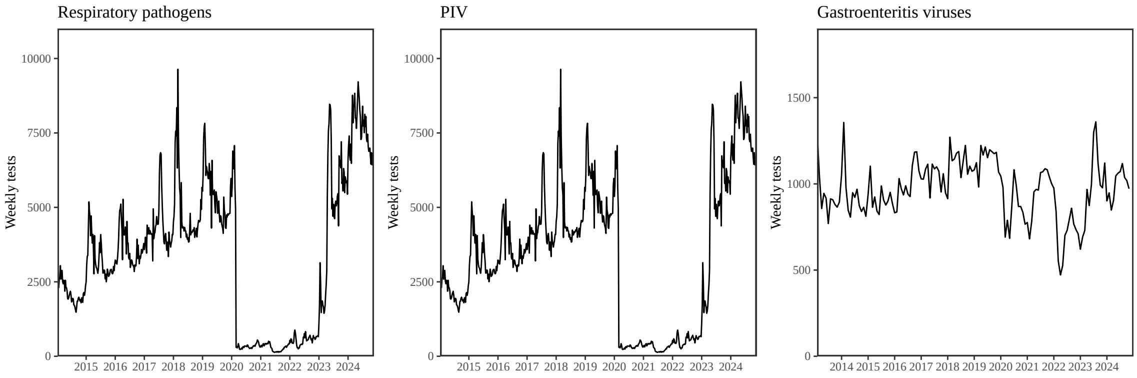


Figure S23: **Testing patterns for respiratory pathogens, PIV, and gastroenteritis viruses in Hong Kong.**

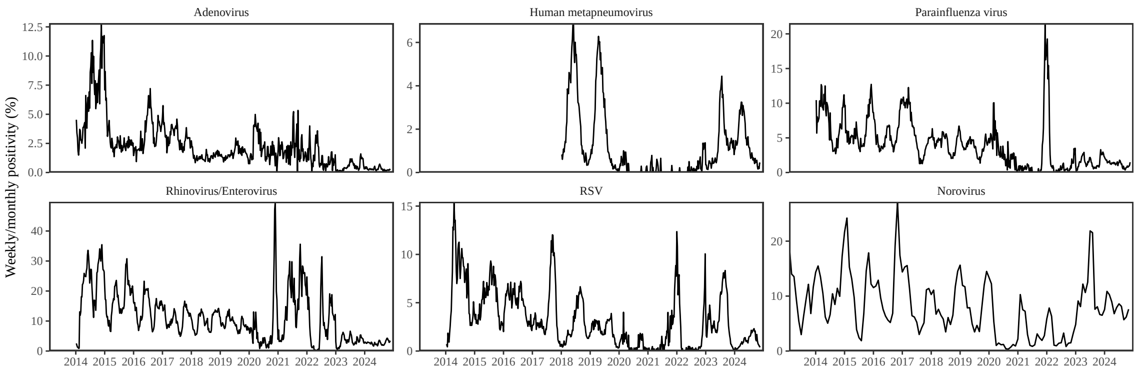


Figure S24: **Test positivity patterns for various pathogens in Hong Kong.**

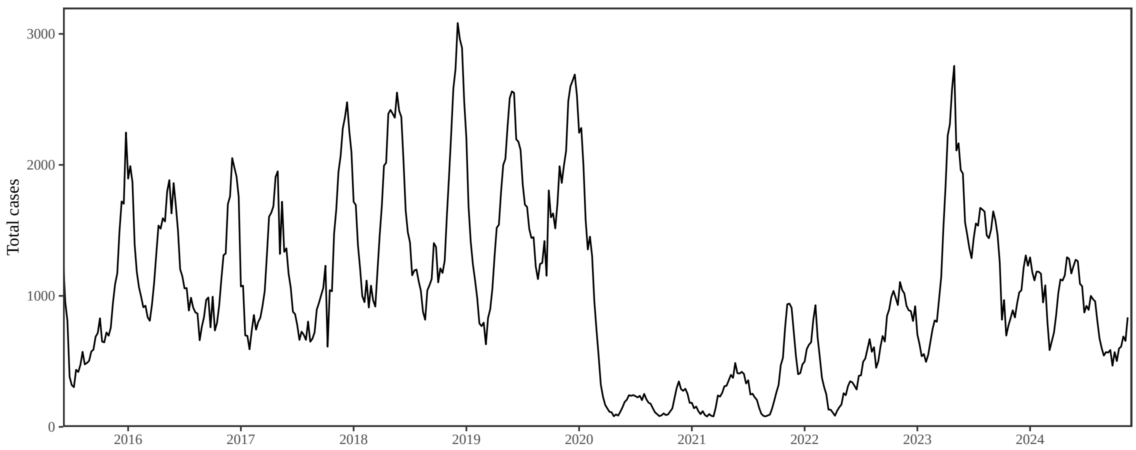


Figure S25: **Total number of reported respiratory infection cases in Korea.**

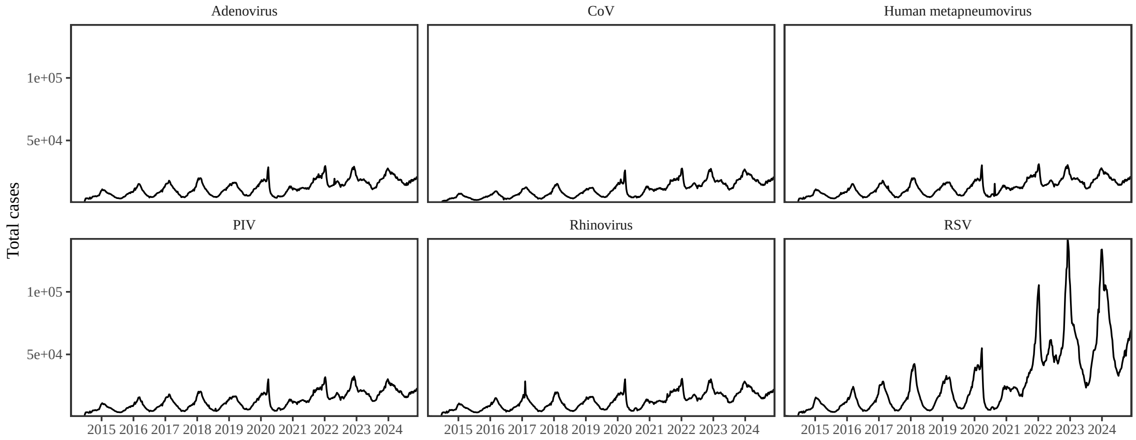


Figure S26: **Testing patterns for respiratory pathogens in the US reported through the NREVSS.**

### References

[1] Virginia E Pitzer, Cécile Viboud, Wladimir J Alonso, Tanya Wilcox, C Jessica Metcalf, Claudia A Steiner, Amber K Haynes, and Bryan T Grenfell. Environmental drivers of the spatiotemporal dynamics of respiratory syncytial virus in the United States. *PLoS pathogens*, 11(1):e1004591, 2015.

[2] Samit Bhattacharyya, Per H Gesteland, Kent Korgenski, Ottar N Bjørnstad, and Frederick R Adler. Cross-immunity between strains explains the dynamical pattern of paramyxoviruses. *Proceedings of the National Academy of Sciences*, 112(43):13396–13400, 2015.

[3] Sang Woo Park, Brooklyn Noble, Emily Howerton, Bjarke F Nielsen, Sarah Lentz, Lilliam Ambroggio, Samuel Dominguez, Kevin Messacar, and Bryan T Grenfell. Predicting the impact of non-pharmaceutical interventions against COVID-19 on Mycoplasma pneumoniae in the United States. *Epidemics*, 49:100808, 2024.

[4] Bjarke Frost Nielsen, Sang Woo Park, Emily Howerton, Olivia Frost Lorentzen, Mogens H Jensen, and Bryan T Grenfell. Complex multiannual cycles of Mycoplasma pneumoniae: persistence and the role of stochasticity. *arXiv*, 2025.

[5] Sang Woo Park, Inga Holmdahl, Emily Howerton, Wenchang Yang, Rachel E Baker, Gabriel A Vecchi, Sarah Cobey, C Jessica E Metcalf, and Bryan T Grenfell. Interplay between climate, childhood mixing, and population-level susceptibility explains a sudden shift in RSV seasonality in Japan. *medRxiv*, pages 2025–03, 2025.

[6] Bob Carpenter, Andrew Gelman, Matthew D Hoffman, Daniel Lee, Ben Goodrich, Michael Betancourt, Marcus A Brubaker, Jiqiang Guo, Peter Li, and Allen Riddell. Stan: A probabilistic programming language. *Journal of statistical software*, 76, 2017.

[7] Stan Development Team. RStan: the R interface to Stan, 2024. R package version 2.32.6.
